# State-Switching Probability Reveals Sleep-Related Biological Drives in *Drosophila*

**DOI:** 10.1101/263301

**Authors:** Timothy D. Wiggin, Patricia R. Goodwin, Nathan C. Donelson, Chang Liu, Kien Trinh, Subhabrata Sanyal, Leslie C. Griffith

## Abstract

Sleep pressure and sleep depth are key regulators of wake and sleep. Current methods of measuring these parameters in *Drosophila melanogaster* have low temporal resolution and/or require disrupting sleep. Here we report a novel analysis tool for high-resolution, non-invasive measurement of sleep pressure and depth from movement data. Probability of transitioning to an active state, P(Wake), measures sleep depth while probability of transitioning to an inactive state, P(Doze), measures sleep pressure. *In vivo* and computational analyses show that P(Wake) and P(Doze) are independent and control the amount of total sleep. Importantly, we demonstrate that these probabilities are tied to specific biological processes. Genetic and environmental perturbations demonstrate that a given amount of sleep can be produced by many combinations of underlying P(Wake) and P(Doze). We show that measuring sleep pressure and depth continuously, without disturbing on-going behavior, provides greater mechanistic insight into behavior than measuring the amount of sleep alone.

## INTRODUCTION

Sleep is a broadly-conserved quiescence behavior that is differentiated from other forms of quiescence (*e.g.* anesthesia, coma) by its internally-driven regulation and fast reversibility (Brown et al. 2010). In the early 20th century, von Economo argued that control of the transition between sleep and wake was localized to a single “nervous center” (von Economo 1930). In contemporary literature, localization of control over sleep and wake is expressed using the idea of “sleep-promoting” or “wake-promoting” structures or cells. However, far from arising from a single center, there are many interconnected cell groups in the mammalian brain that participate in wake/sleep transitions (Scammell et al. 2017). Even in the fruit fly *Drosophila melanogaster*, which has a much smaller brain, the localization of sleep initiation is complex, with dozens of cellular loci that can drive sleep or activity (Tomita et al. 2017). In addition to neuronal mechanisms, there is a rich pallet of hormonal and metabolic factors that tip the balance between sleep and wake (Nässel and Zandawala 2019; Yurgel et al. 2015). There are not currently tools for systematically understanding how the many biological drives toward wake and sleep are integrated, especially how they combine with, synergize with, or occlude one another.

The difficulty in integrating these different signals may arise from focusing on a single behavioral measure, the amount of sleep, rather than capturing biological drives that regulate the amount of sleep. In humans, sleep is regulated both by the intensity of the drive to fall asleep (higher in narcolepsy, lower in insomnia) and the tendency to wake up (higher in insomnia, lower in hypersomnolence and depression) (Barateau et al. 2017; Kales and Kales 1974). To capture these drives, conditional probability models using neuronal firing or EEG data have been used to model the structure of healthy and disordered sleep in mammals (Bianchi et al. 2012; Perez-Atencio et al. 2018; Stephenson et al. 2016; Yang and Hursch 1973), but this approach had not yet been applied to sleep in *Drosophila*.

In this paper, we use conditional probability of state switching to measure the biological drives of sleep in *Drosophila*. We define two conditional probabilities: P(Doze), the probability that an active fly will stop moving, and P(Wake) the probability that a stationary fly will start moving. Using an *in silico* model of sleeping flies, we demonstrate that the combination of P(Doze) and P(Wake) is sufficient to explain the total amount of time flies spend asleep. We experimentally determine that P(Doze) and P(Wake) are measures of sleep pressure and sleep depth, respectively. We find that P(Wake) is strongly influenced by dopamine, the major neurochemical involved in arousal in *Drosophila* (Andretic et al. 2005; Kume et al. 2005), and that sleep structure can be regulated by P(Doze) as well as P(Wake). Finally, we find that age-dependent changes in sleep (Koh et al. 2006; Vienne et al. 2016) reflect changes in the balance of P(Doze) and P(Wake) and that measurement of transition probabilities reveals novel aging/sleep interactions.

## RESULTS

### Conditional Wake and Doze Probability Determine Drosophila Sleep

Measurement of the amount of sleep in *Drosophila* is based on movement data, typically taken in bins of ≤ 1 min, where a sleep episode is defined as certain period of time without movement, usually 5 min (Hendricks et al. 2000; Shaw et al. 2000).

Between each individual observation, the fly may choose to either remain in its current state or switch. The sequence of states reveals not only the amount of time spent in each state (*i.e.* time asleep), but also the conditional probability of switching states *e.g*. from active to inactive. We call these behavioral transition probabilities P(Doze), the probability that the fly will switch from the active state to the inactive state, and P(Wake), the probability that the fly will switch from the inactive state to the active state, (Fig. 1A, see *Methods* for detailed explanation).

**Figure 1.**
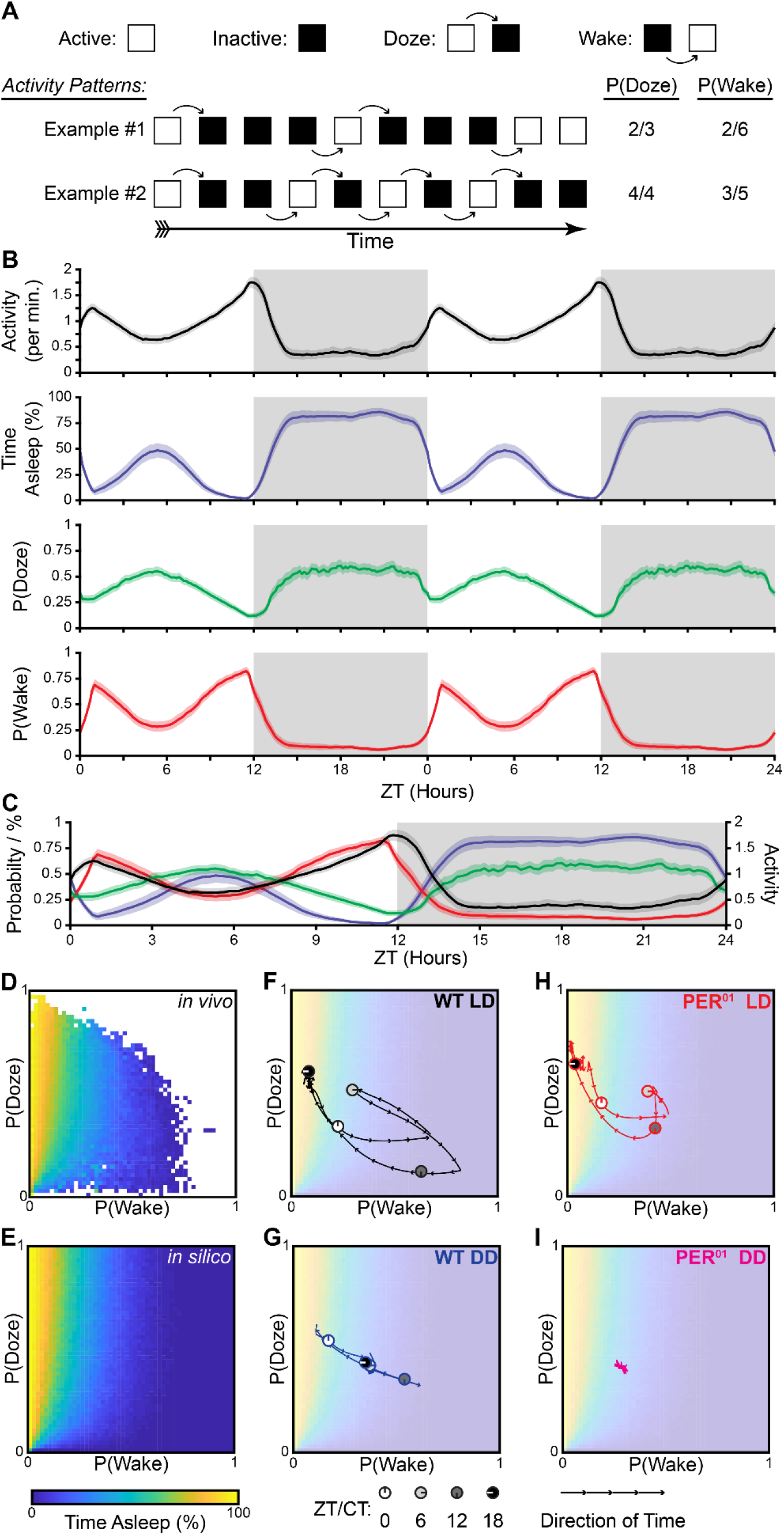
Conditional Wake and Doze Probability Determine Drosophila Sleep. (A) Two exemplar activity patterns (Examples # 1 & 2). White squares: wake, Black squares: sleep. Transitions between behavioral states are indicated by curved arrows. Transitions from active to inactive (Doze) are above the blocks, transitions from inactive to active (Wake) are below. Calculation of P(Wake) and P(Doze) are described in *Methods*. (B) Double plotted daily cycles of Activity, % Time Asleep, P(Doze), and P(Wake) measured in WT female flies. Solid lines are the population mean, shaded area is the 95% confidence interval of the mean. Individual profiles are the average of 3 days of behavior. Gray area indicates lights off period. ZT = Zeitgeber Time, or hours since lights on. (C) The same data presented in (B) plotted on a common set of axes. Color coding is the same as (B). (D-E) Heatmaps of *in vivo* (D) and *in silico* (E) sleep properties. The behavior of each *in vivo* fly was split into 90 minute time intervals, and the P(Wake), P(Doze), and total sleep for that interval was calculated. Each 90 minute behavioral sample was assigned to one of 51 bins from 0 to 1 based upon its respective P(Doze) and P(Wake). The mean value of total sleep was calculated for the samples in each bin, and the bin was assigned a color according to the legend (*bottom*). Coordinates without any biological samples are displayed as white. (E) For each P(Doze) and P(Wake) bin, 64 *in silico* flies were simulated. The mean total sleep in these simulations is displayed using the same color map as the *in vivo* data. (F-I) Plots of the mean trajectory of flies through the P(Wake) vs. P(Doze) probability space across their daily rhythm superimposed on the *in silico* predicted total sleep at each point. The direction of time is indicated by arrowheads. Important circadian times are indicated by small clock faces (see key at *bottom*). Genotype and environmental conditions are indicated in the corner of each plot. PER^01^: *period* point mutation, LD: 12:12 light:dark cycle, DD: constant darkness.

These probabilities should change in response to changes in sleep drive and arousal. In order to assess the pattern of behavioral state transition probability, we analyzed behavioral data acquired with both the well-characterized *Drosophila* Activity Monitor (DAM) system (*Canton Special* (WT) female flies; *n =* 60; Fig. 1) and with the more recently developed FlyBox (WT female flies; *n =* 55; Fig. S1) (Guo et al. 2016). P(Wake) and P(Doze) both change across the day, but do not exactly replicate activity, sleep, or one another regardless of the method of data acquisition (Fig. 1B-C, S1C). Both P(Wake) and P(Doze) have strong circadian influences, with P(Wake) showing a similarity to locomotor activity. P(Doze) cycles more weakly and circumscribes sleep.

In order to assess the degree of interdependence among these variables quantitatively, we measured the between-animal Pearson correlation of the measures. P(Wake) and total sleep are strongly anticorrelated (*R* = −0.92, *p* < 0.0001), while P(Doze) and total sleep are correlated, but less strongly (*R* = 0.26, *p* < 0.0001). Interestingly, P(Wake) and P(Doze) are only moderately anticorrelated with one another (*R* = −0.18, *p* < 0.0001). These relationships, and the shape of the curves in Fig. 1B, suggest that the transition probabilities are not simply each other’s inverse, but are instead likely to be driven by distinct biological mechanisms.

The three-way relationship between P(Wake), P(Doze), and sleep can be illustrated using a heatmap of total sleep generated at each combination of probabilities (Fig. 1D). In order to interrogate the causal relationships between these variables, we used a Markov-chain model to generate *in silico* behavior. For the probability bins in which we have *in vivo* data, there is an extremely good match between the *in vivo* and *in silico* data (RMSE = 5.8% Time Asleep, *R* = 0.99, *p* < 0.0001). The lack of unexplained variance in the *in vivo* data with respect to the *in silico* data suggests strongly that the combination of P(Wake) and P(Doze) causally determines the amount of total sleep for each individual fly.

The daily behavioral cycle of flies can be visualized as trajectories through this probability space (Fig. 1F-I). In wild type animals, the presence of a light cycle generates a double “U”-shaped trajectory (Fig. F). The shape reflects movement of animals from high sleep regions of the probability space at night (ZT18) and during siesta (ZT6) to their most active phases at the light/dark transitions (ZT0 and ZT12). Each time of day is associated with a different balance of P(Wake) and P(Doze).

In contrast to behavior in light:dark cycles, wild type flies in constant darkness (Fig. 1 G) have a purely linear trajectory through probability space that is largely due to changes in P(Wake), with P(Doze) showing little change (Fig. 1G). It is notable that the relative positions in probability space of animals at the circadian times when light:dark transitions are expected (CT0 and CT12) appear to be similar to those of animals in cycling conditions, suggesting that at these times of day P(Wake) and P(Doze) may be purely clock-driven. The P(Wake)/P(Doze) position of animals at times when they are normally asleep (ZT6 and ZT18) are strongly affected by light cycles, which increases the amount of sleep at both times. The differences in position in probability space between LD and DD suggest that siesta sleep is increased by a light using a mechanism involving only changes in P(Doze), while the light-dependent enhancement of nighttime sleep is due to movement along both the P(Doze) and P(Wake) axes.

To test the role of the circadian clock in specification of transition probabilities, we carried out the same analyses on animals lacking a functional clock (Fig. 1 H and I). In cycling light, these animals showed a U-shaped trajectory similar to wild type, indicating that the divergence from linearity caused by light does not require an intact clock. In the absence of both light cues and a circadian clock, flies simply find their behavioral set point and remain stationary in the probability space. This implies that the clock controls the light-independent regulation of both P(Wake) and P(Doze). Given the apparent importance of P(Wake) and P(Doze) in setting the level of total sleep, we sought to relate each to a biological process.

### Conditional Wake Probability is a Measure of Sleep Depth

Sleep is a state in which movement and responses to external stimuli are suppressed. The more strongly these behaviors are suppressed, the “deeper” the sleep. Because the likelihood of waking up is logically linked to the depth of sleep, we hypothesized that P(Wake) is a measure of sleep depth. Sleep depth is most commonly measured in flies using arousal threshold (van Alphen et al. 2013; Guo et al. 2011; Hendricks et al. 2000; Huber et al. 2004; Linford et al. 2012; Shaw et al. 2000). To examine the relationship between transition probabilities and arousal state, we measured P(Wake), P(Doze), and arousal threshold at two temperatures: a standard temperature (25°C) and elevated temperature (29-30°C). Since elevated temperature increases sleep during the day and decreases it at night (Parisky et al. 2016), we hypothesized that sleep depth should also be affected by the change in temperature. As predicted, temperature affects both P(Wake) and P(Doze). P(Wake) is significantly decreased during the daytime (*p* < 0.0001) and elevated during the nighttime (*p* < 0.0001) at 29°C (Fig. 2B). Daytime P(Doze) is significantly elevated by heat (*p* < 0.0001), but nighttime P(Doze) is unchanged (*p* = 0.057).

**Figure 2.**
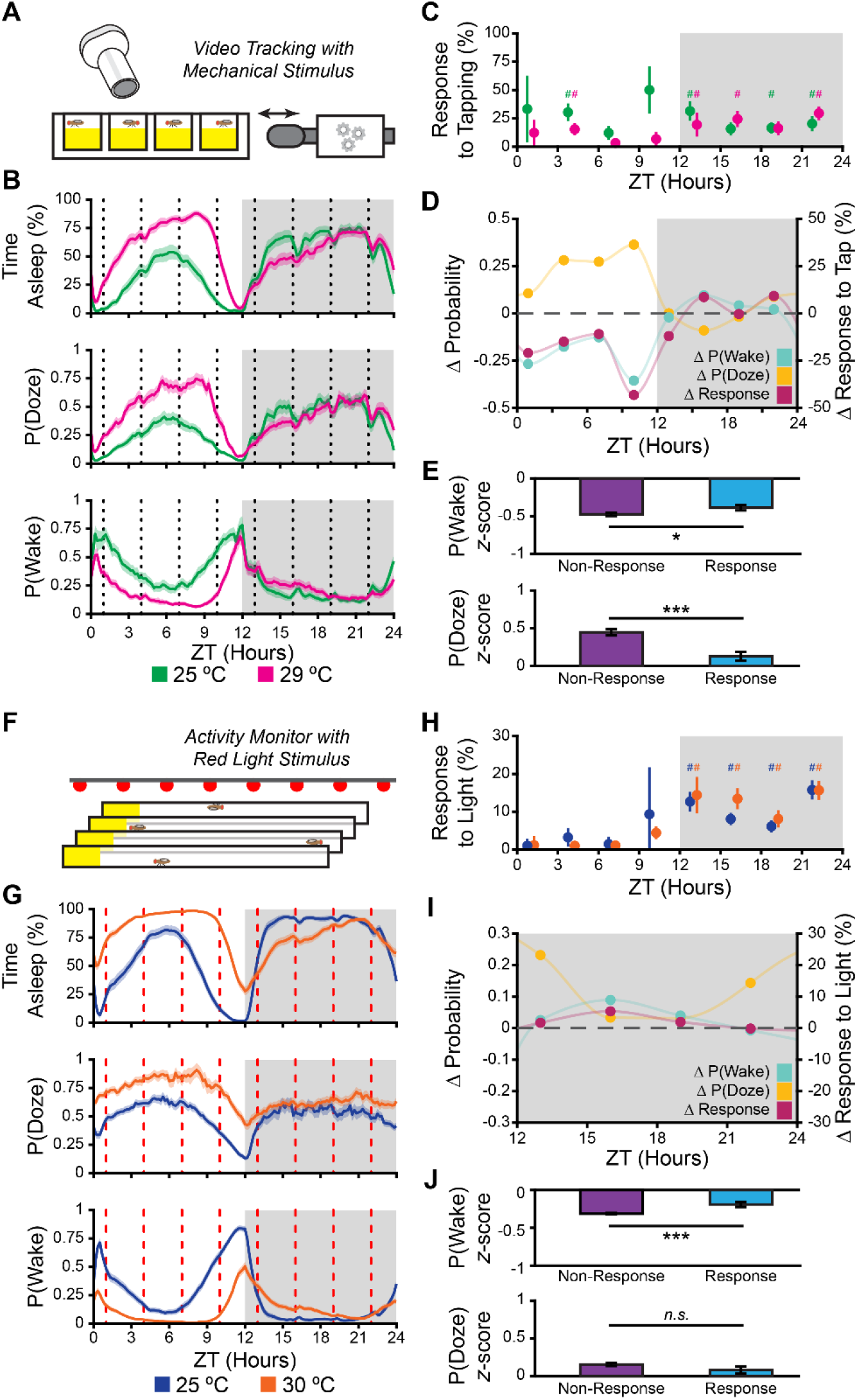
Conditional Wake Probability is a Measure of Sleep Depth. (A-E) Results of a mechanical arousal threshold experiment. (A) Schematic of experimental set-up. Fly locomotion in a 96-well plate was monitored using a FlyBox. At 3 hour intervals starting at ZT1, an electromagnetic solenoid was used to tap the side of the plate. (B) Circadian profiles of % Time Asleep, P(Doze), and P(Wake). Solid lines are the population mean, shaded area is the 95% confidence interval. Behavior was measured at 25°C *(green)* and 29°C *(magenta)*. Tapping times are indicated by dashed vertical lines. Gray area indicates lights off period. (C) Response to tapping was measured as the increase in arousal probability following a tap. The % responding is plotted at 25°C *(green)* and 29°C *(magenta)*. # indicates response greater than chance (*p* < 0.05) measured by Fisher’s exact test with the Holm’s step-down correction for multiple testing. (D) The effect of temperature on P(Wake) *(cyan)*, P(Doze) *(yellow)*, and response to tapping *(pink)* at each time (ZT). (E) The P(Wake) and P(Doze) of each timepoint was normalized by converting it to a *z*-score. The behavior of each fly following each tap was classified as a non-response (no activity 1 minute after tap) or response. The mean *z*-score across all time points for responders (cyan bars) and non-responders (purple bars) is plotted. (F-J) Results of a light arousal threshold experiment. (F) Schematic of experimental set-up. Fly locomotion in glass tubes was monitored using the DAM system. At 3 hour intervals starting at ZT1, an array of red LEDs was flashed for 2 seconds. (G) Circadian profiles of % Time Asleep, P(Doze), and P(Wake). Solid lines are the population mean, shaded area is the 95% confidence interval. Behavior was measured at 25°C *(blue)* and 30°C *(orange)*. LED flash times are indicated by dashed vertical lines (*red*). (H) Response to light was measured as the increase in arousal probability following a light flash, as in (C). The % responding is plotted at 25°C *(blue)* and 29°C *(orange)*. (I) The effect of temperature on P(Wake) *(cyan)*, P(Doze) *(yellow)*, and response to light *(pink)* at each time (ZT). (J) P(Wake) and P(Doze) were normalized, and the behavior of each fly following each light flash was classified, as in (E). The normalized P(Wake) and P(Doze) for non-response and response. Error bars are standard error of the mean. *n.s.* indicates no significant difference, * p < 0.05, ** p < 0.005, *** p < 0.0005.

Because sensitivity to sensory stimuli is also regulated by time of day (van Alphen et al. 2013; Chen et al. 1992; Krishnan et al. 1999), we measured mechanical arousal threshold with gentle tapping at 3-hour intervals across the day using FlyBox (ZT 1, 4, 7, 10, 13, 16, 19, 22; *n* = 89 WT female flies; Fig. 2A). Tapping the flies produced an excess of awakenings compared to chance at timepoints across the daily cycle at both baseline and high temperatures (Fig. 2C). Heat produces significantly anticorrelated effects on P(Wake) and P(Doze) (*R* = −0.82, *p* = 0.012), so the method of partial correlations was used to determine their independent associations with arousal threshold. The effect of heat on the response to tapping across time points is significantly correlated the effect of heat on P(Wake) (*R* = 0.84, *p* = 0.016) but not the effect of heat on P(Doze) (*p* = 0.77; Fig. 2D). Normalizing P(Wake) and P(Doze) between timepoints, individual flies that responded to the tap had a significantly higher P(Wake) and significantly lower P(Doze) than the non-responsive flies (*p* = 0.024 and p < 0.0001, respectively; Fig. 2E). We conclude from this experiment that P(Wake) and arousal threshold are regulated identically by heat, in line with our prediction.

As an independent probe of the association of transition probabilities with sleep depth, we measured light-mediated arousal threshold with flashes of red light across the day using the DAM system (*n* = 117 WT female flies; Fig. 2F). As we found in the FlyBox, increasing temperature decreases P(Wake) during daytime and increases it during nighttime (*p* < 0.0001). In contrast to the mechanical arousal experiment, in this experiment we found that P(Doze) was significantly elevated during both daytime and nighttime (*p* < 0.0001). Unlike mechanical tapping, light flashes were only sufficient to rouse the flies during the lights-off period (Fig. 2H), so we confined our analyses of arousal threshold to these times of day. Comparing across timepoints, P(Doze) and P(Wake) are not significantly correlated (p = 0.07), so Pearson correlation was used to measure the association between arousal probability and each transition probability. The effect of heat on light arousal is highly correlated with the effect of heat on P(Wake) (*R* = 0.92, *p* = 0.02) but not with the effect of heat on P(Doze) (*p* = 0.06). Within each timepoint, individual flies that responded to the light had a significantly higher P(Wake) than the non-responsive flies (*p* < 0.0001) but there was no difference in P(Doze) (*p* = 0.15; Fig. 2J).

In summary, the effect of heat on arousal threshold is tightly coupled with the effect of heat on P(Wake) regardless of sensory modality or monitoring system used. Coupling between mechanical arousal threshold and P(Doze) was only seen when normalizing P(Doze) between time points in the comparison of responding vs. non-responding individuals (Fig. 2E). We posit that the association in this experiment is due to the latent correlation between P(Wake) and P(Doze) noted previously and does not reflect a true correlation of arousal threshold with P(Doze). This latent correlation is removed by the partial linear correlation method (used to analyze data in Fig. 2D where there is no P(Doze) / arousal threshold association) but is not removed in the comparison of responders and non-responders in Fig. 2E. We therefore conclude that P(Wake) and arousal threshold, but not P(Doze), measure the same underlying biological process, namely, sleep depth.

### Conditional Doze Probability is a Measure of Sleep Pressure

Sleep deprivation in flies, like in other animals, leads to increased subsequent sleep (Hendricks et al. 2000; Shaw et al. 2000). A common model for the generation of increased sleep after release from sleep deprivation posits that there is an increase in sleep “pressure” which increases sleep drive (Borbély 1982; Fuller et al. 2006). Because sleep drive increases the tendency to fall asleep, we hypothesized that P(Doze) is related to sleep pressure.

To increase sleep pressure and examine its relationship to P(Doze), we performed a nighttime sleep deprivation (SD) by shaking flies between ZT12-24 (WT female flies; *n* = 120 SD, 117 Control; Fig. 3A). The experimental flies sleep less during SD than they do when unperturbed, acquiring sleep debt which is discharged when they are released from SD (Fig. 3B). While flies are being shaken, P(Wake) is increased and P(Doze) is decreased (Fig. 3C). After the flies are released from SD, there is an immediate, significant increase in P(Doze) and a decrease in P(Wake) (ZT0-6; *p* < 0.0001; Fig. 3D,E). The effect on P(Doze) disappears after the initial recovery morning (ZT0-6), while there is a residual effect of sleep deprivation on P(Wake) in early night sleep (ZT12-18; *p* = 0.002; Fig. 3E). In this experiment both P(Wake) and P(Doze) are altered by SD and neither can be excluded as a measure of sleep pressure.

**Figure 3.**
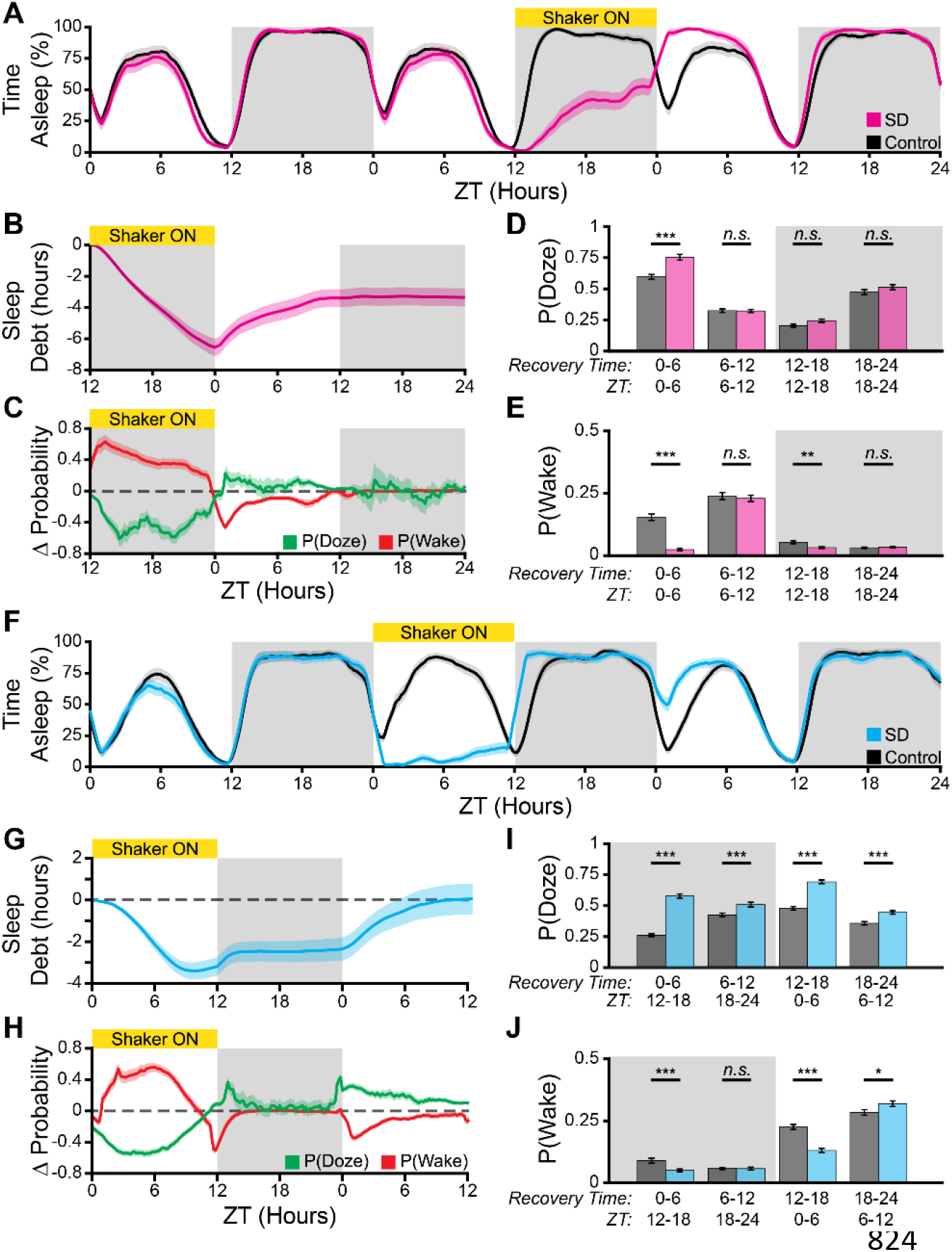
Conditional Doze Probability is a Measure of Sleep Pressure. (A-E) Results of a nighttime sleep deprivation experiment. (A) % Time asleep of shaken *(magenta)* and unshaken control *(grey)* flies over 3 days, including one night of sleep deprivation (labeled “Shaker ON”). Shaded regions indicate lights on and lights off times. (B) 824 Sleep debt, cumulative sleep minus baseline sleep, accumulated during sleep deprivation and recovered following release from deprivation. (C) Change in P(Doze) *(green)* and P(Wake) *(red)* during sleep deprivation and following release from deprivation. (D,E) Comparison of P(Doze) (D) and P(Wake) (E) between shaken *(magenta)* and control *(grey)* flies following release from sleep deprivation. (F-J) Results of a daytime sleep deprivation experiment. (F) % Time asleep of shaken *(cyan)* and unshaken control *(grey)* flies over 3 days, including one day of sleep deprivation (G) Sleep debt accumulated during daytime sleep deprivation and recovered following release from deprivation. (H) Change in P(Doze) *(green)* and P(Wake) *(red)* during sleep deprivation and following release from deprivation. (I,J) Comparison of P(Doze) (I) and P(Wake) (J) between shaken *(cyan)* and control *(grey)* flies following release from sleep deprivation. Error bars are standard error of the mean. *n.s.* indicates no significant difference, * p < 0.05, ** p < 0.005, *** p < 0.0005.

We next performed a daytime (ZT0-12) sleep deprivation experiment (WT female flies; *n* = 116 SD, 118 Control; Fig. 3F). Because the expression of recovery sleep is regulated by time of day (Cavanaugh et al. 2016), we hypothesized that the sleep homeostat would maintain a “memory trace” of sleep debt during times of day when recovery sleep could not be expressed. As in the nighttime sleep deprivation experiment, flies acquire sleep debt during shaking, P(Wake) is upregulated, and P(Doze) is down regulated (Fig. 3G,H). Following release from sleep deprivation at ZT12, P(Doze) is upregulated, but sleep debt is only minimally discharged during the night, with most of the recovery sleep occurring during the next light period (Fig. 3G). Unlike the situation with nighttime SD, P(Doze) remains elevated for 24 hours until sleep debt is fully discharged (p < 0.0005; Fig. 3I). In contrast, P(Wake) is significantly suppressed only during the times of day when recovery sleep is actually being performed, *i.e.* in the early evening (ZT12-18; *p* = 0.0005) and in the morning pre-siesta (ZT0-6; *p* < 0.0001; Fig. 3J).

These experiments demonstrate that P(Doze) has properties consistent with a measure of sleep pressure: it is elevated by sleep deprivation and returns to baseline only after sleep debt is discharged. In contrast, suppression of P(Wake) is necessary for the expression of rebound sleep but does not have the same memory trace properties as P(Doze). Interestingly, however, P(Doze) also appears to be regulated by “wake pressure” in addition to sleep pressure. This is most obvious during the shaking period, when P(Doze) is down-regulated because of disruptive mechanosensory inputs provide an environmentally-driven wake pressure (Fig. 3G,H). There are also likely effects of internally-generated wake pressure. These can also be inferred from the decrease in P(Doze) during the anticipatory evening activity peak before ZT12 in unperturbed flies (Fig. 1B). We conclude that P(Doze) is a measure of sleep pressure, with the caveat that strongly wake-maintaining processes (sensory, circadian, or otherwise) can mask our ability to detect sleep pressure via behavioral outputs.

### Dopaminergic Tone Regulates Sleep Depth

Sleep is regulated by a multitude of molecular processes, and many null and hypomorphic mutations disrupt sleep in flies (Helfrich-Förster 2018). Dopamine in particular has a role in regulating sleep quantity by increasing arousal (Andretic et al. 2005; Kume et al. 2005) and acute activation of dopaminergic neurons has been shown to decrease total sleep (Seidner et al. 2015; Shang et al. 2011; Ueno et al. 2012b; Vienne et al. 2016). In order to validate the proposed biological significance of P(Wake) and P(Doze) described above, we measured the effect of manipulating dopaminergic tone on behavioral state transition probability by performing a 2-day thermogenetic activation (Hamada et al. 2008) of *TH-Gal4* positive dopamine neurons (Fig. 4A). In order to separate the effect of dopamine neuron activation from the direct effect of temperature on sleep (Fig. 2), we compared the effect of our heat manipulation on the experimental line (*TH-Gal4/UAS-dTrpA* female flies, *n* = 31) with the effect on parental strain controls (*TH-Gal4/+* female flies, *n* = 31; *UAS-dTrpA*/+ female flies, *n* = 61). An effect on behavior was only attributed to dopamine neuron activation when the experimental line was significantly different from both control genotypes.

**Figure 4.**
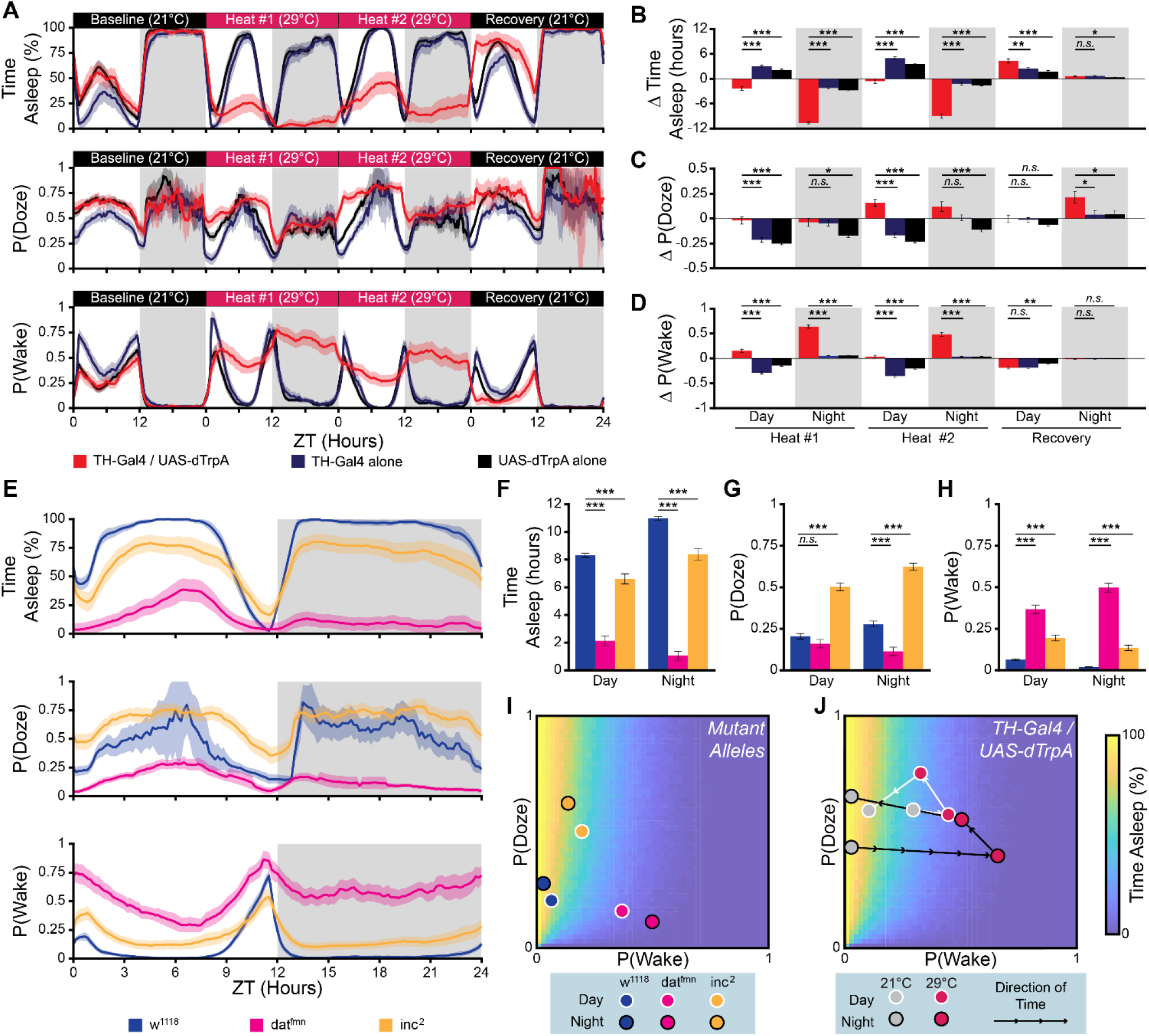
Dopaminergic Tone Regulates Sleep Depth. (A) Sleep, P(Doze), and P(Wake) during the thermogenetic activation of dopamine neurons. The experimental flies *(red)* were compared with control flies carrying only the driver *(TH-Gal4 alone, blue)* or only the actuator *(UAS-dTrpA, black)*. (B-D) Comparison of the heat-induced change in total sleep (B), P(Doze) (C), and P(Wake) (D) between experimental flies and control flies. (E) Circadian profile of % time asleep, P(Doze), and P(Wake) for *dat*^*fmn*^ (*magenta*), *inc*^2^ (*yellow*), and control flies (*w*^*1118*^, *blue*). (F-H) Comparison of total sleep (F), P(Doze) (G), and P(Wake) (H) between the experimental flies and the control flies. (I) The population mean P(Wake) and P(Doze) for each mutant line and the control flies is overlaid on the *in silico* predicted sleep heatmap. (J) The population mean of the *TH-Gal4/UAS-dTrpA* experimental group during the thermogenetic activation experiment is overlaid on the *in silico* predicted sleep heatmap. Arrows indicate the direction of time, grey points: 21°C, red points: 29°C. In (I) and (J), circles with white borders are daytime values, circles with black borders are nighttime values. Error bars are standard error of the mean. *n.s.* indicates no significant difference, * p < 0.05, ** p < 0.005, *** p < 0.0005.

As has been previously shown, activating dopamine neurons significantly reduces total sleep throughout the activation period (*p* < 0.0001; Fig. 4B). This is accompanied by a very substantial increase in P(Wake) throughout the activation period (*p* < 0.0001; Fig. 4D) with only a modest daytime increase in P(Doze) (*p* < 0.0001; Fig. 4C). Following the cessation of activation, the experimental line had significantly increased sleep (*p* < 0.0018), but P(Wake) and P(Doze) were not significantly altered compared to controls. Close examination of the behavioral traces (Fig. 4A) suggests that there may be transient differences in P(Wake) and P(Doze) immediately after temperature is restored to baseline, but the change is not maintained long enough to reach statistical significance when averaged across the entire lights-on period.

Because temperature has a strong effect on sleep independent of dopaminergic activation, we also investigated chronic genetic perturbations of the dopamine system. We examined the effects of two dopamine-associated sleep mutants, *fumin* (*dat*^*fmn*^ male flies; n = 37) (Kume et al. 2005) and *insomniac* (*inc*^2^ male flies; n = 31) (Stavropoulos and Young 2011). We compared the behavioral state transition probabilities of these mutants with control flies of the same genetic background (*w*^1118^ male flies; *n* = 40; Fig. 4E). Both *fumin* and *insomniac* flies have decreased total sleep (*p* < 0.0002; Fig.4F), and increased P(Wake) (*p* < 0.0001; Fig. 4H). However, these mutants differ in their effect on P(Doze): *insomniac* significantly increases P(Doze) (*p* < 0.0001), while *fumin* decreases P(Doze), but the effect is only statistically significant during the night (*p* < 0.0001; Fig. 4G). Both acute activation of dopamine neurons and chronic increases of dopaminergic tone clearly decrease sleep and increase P(Wake), supporting the idea that P(Wake) is a measure of sleep depth. The effect of dopamine on P(Doze) is less consistent and is dependent on genetic context.

Plotting the mutant fly data in probability space shows that the somewhat divergent total sleep phenotypes of these mutants are predicted by our *in silico* simulation of the P(Wake)/P(Doze) relationship to sleep amount (Fig. 4I). The decrease in total sleep compared to wild type is clearly attributable to the change in P(Wake) for both mutants. *insomniac* mutants have a P(Doze) set point that is very distinct from either wild type or *fumin* animals, but both mutants have normal day-to-night shifts in P(Wake)/P(Doze), suggesting that the clock-mediated process that regulate this difference are intact.

The total sleep effects of acute changes in dopaminergic neuron activity are also predicted by our *in silico* model (Fig. 4J). Interestingly, the first 24 h of acute dopamine neuron activation (trajectories from first gray circle to first red circle) resembles the *fumin* phenotype: dramatically increased P(Wake) and little to no P(Doze) effect. The second 24 h of activation (trajectories from first red circle to second red circle) more closely resembles *insomniac*: less dramatically increased P(Wake) compared to baseline accompanied by increased P(Doze). This may be indicative of underlying differences between the mechanisms of *insomniac* and *fumin* in regulating sleep and supports the idea that *insomniac* affects homeostatic processes (Pfeiffenberger and Allada 2012).

### High Sugar Diet Promotes Sleep Consolidation by Decreasing Sleep Pressure

Both sleep deprivation and manipulations of the dopamine system produce dramatic changes in total sleep (Fig. 3 and 4). In order to test the relationship of P(Wake) and P(Doze) to sleep structure, we measured behavior of WT flies fed either a low (2.5%) or high (30%) concentration of sucrose, a manipulation which has been shown to regulate sleep fragmentation independent of total sleep (Linford et al. 2012). In agreement with these findings, we do not observe dramatic changes in the daily sleep pattern between low and high sucrose diet (WT female flies; *n* = 90 low sucrose, 94 high sucrose; Fig. 5A). The effects on the amount of sleep are modest: no effect of diet on daytime sleep, and a small increase in nighttime sleep (*p* = 0.0001; Fig. 5B).

**Figure 5.**
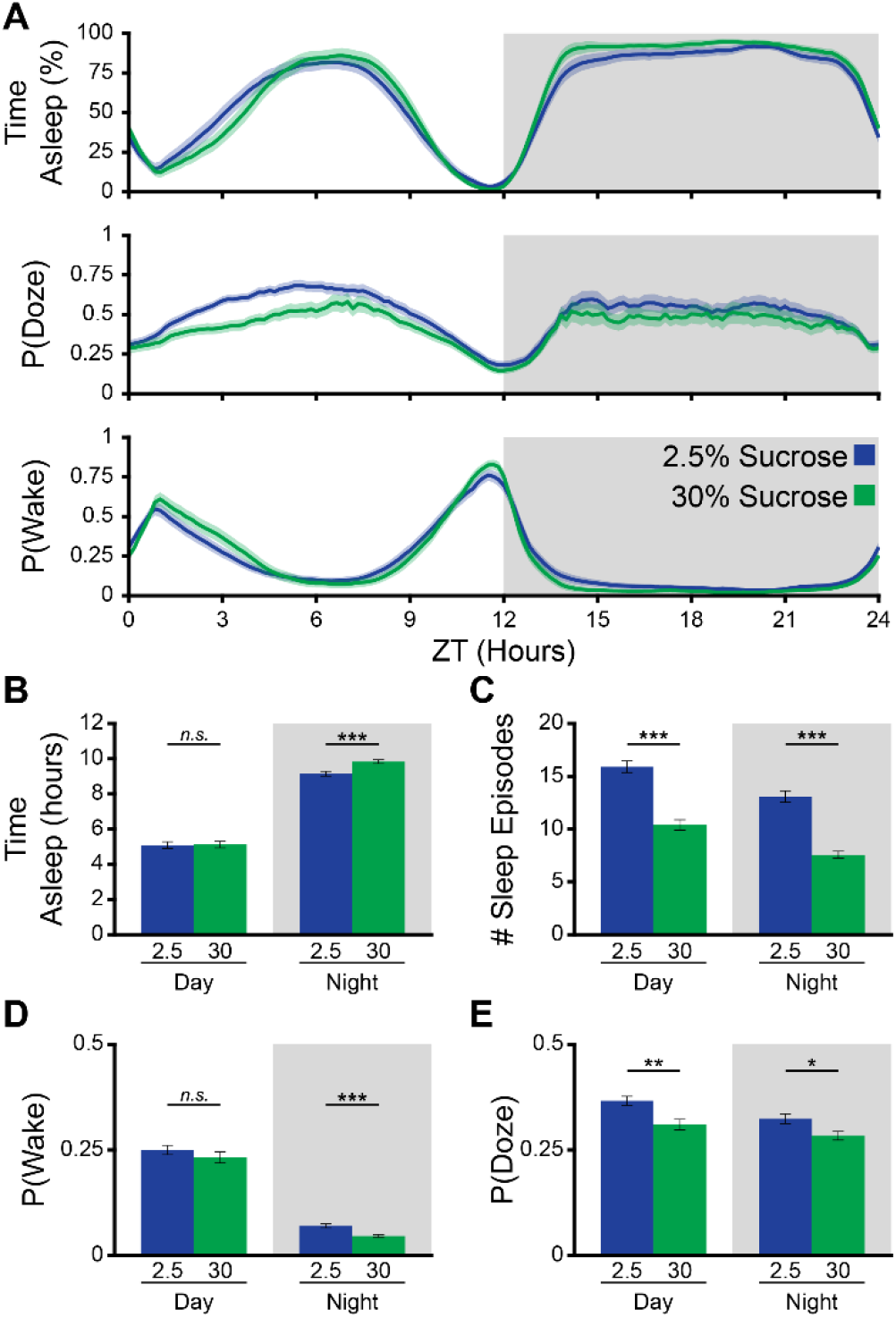
High Sugar Diet Promotes Sleep Consolidation by Decreasing Sleep Pressure. (A) Daily cycle of % time asleep, P(Doze), and P(Wake) for wildtype (CS) flies fed either a low sugar (2.5% sucrose in agar, blue), or a high sugar (30% sucrose in agar, green) diet. (B-E) Comparison of low and high sugar diet time asleep (B), # sleep episodes (C), P(Wake) (D), and P(Doze) (E) during daytime and nighttime. *n.s.* indicates no significant difference, * p < 0.05, ** p < 0.005, *** p < 0.0005.

There is, however, a large decrease in the number of sleep episodes both during the day and at night in the high sucrose group (*p* < 0.0001; Fig. 5C). Interestingly, we find that P(Wake) is unaffected by diet during the daytime but significantly decreased at night (*p* < 0.0001; Fig. 5D), in concordance with the effect of sucrose on total sleep. P(Doze) is decreased by high sucrose both during daytime and nighttime, mirroring the effect of sucrose on sleep structure (*p* < 0.009; Fig. 5E). This was an unexpected finding, because the initial report of a sucrose effect on sleep structure attributed the effect to sleep depth (Linford et al. 2012). This discrepancy is likely due to this prior study measuring sleep depth (via arousal threshold) only at night, a time at which we also find a small difference in P(Wake) (Fig. 5D). With our improved analysis tools, sleep depth and pressure can be estimated independently throughout the day, allowing us to separate the different dimensions of diet/environmental interactions and conclude that the change in sleep structure is more likely due to P(Doze), a surprising finding since fragmentation is usually thought to be due to changes in arousal (see Discussion).

### Changes in Sleep with Age are Due to Both Sleep Pressure and Depth

To test the utility of our analysis tools in understanding a complex, longitudinal biological process, we measured the behavior of flies across the lifespan. The total amount and structure of sleep produced by an individual is strongly regulated by age in many animals, including flies (Koh et al. 2006; Vienne et al. 2016). Sleep was measured in cohorts of WT flies during week 1, 3, 5, and 7/8 post eclosion (*n* = 48, 30, 30, and 39, respectively; weeks 7 & 8 were combined due to a low rate of survival to this age). In agreement with prior results, the daily profile of sleep is altered in aged flies, most dramatically during the day (Fig. 6A). In these cohorts of flies, old flies slept significantly longer than young flies (p < 0.0002; Fig. 6B). Perhaps surprisingly given our results with groups of young flies (Fig. 1-3) where the amount of sleep was determined largely by changes in P(Wake), nighttime P(Wake) does not vary significantly as flies age, and daytime P(Wake) is only significantly decreased in the oldest group of flies (p < 0.0001; Fig. 5D). In contrast, P(Doze) increases with age both during the day and night (p < 0.0001; Fig. 5C).

**Figure 6.**
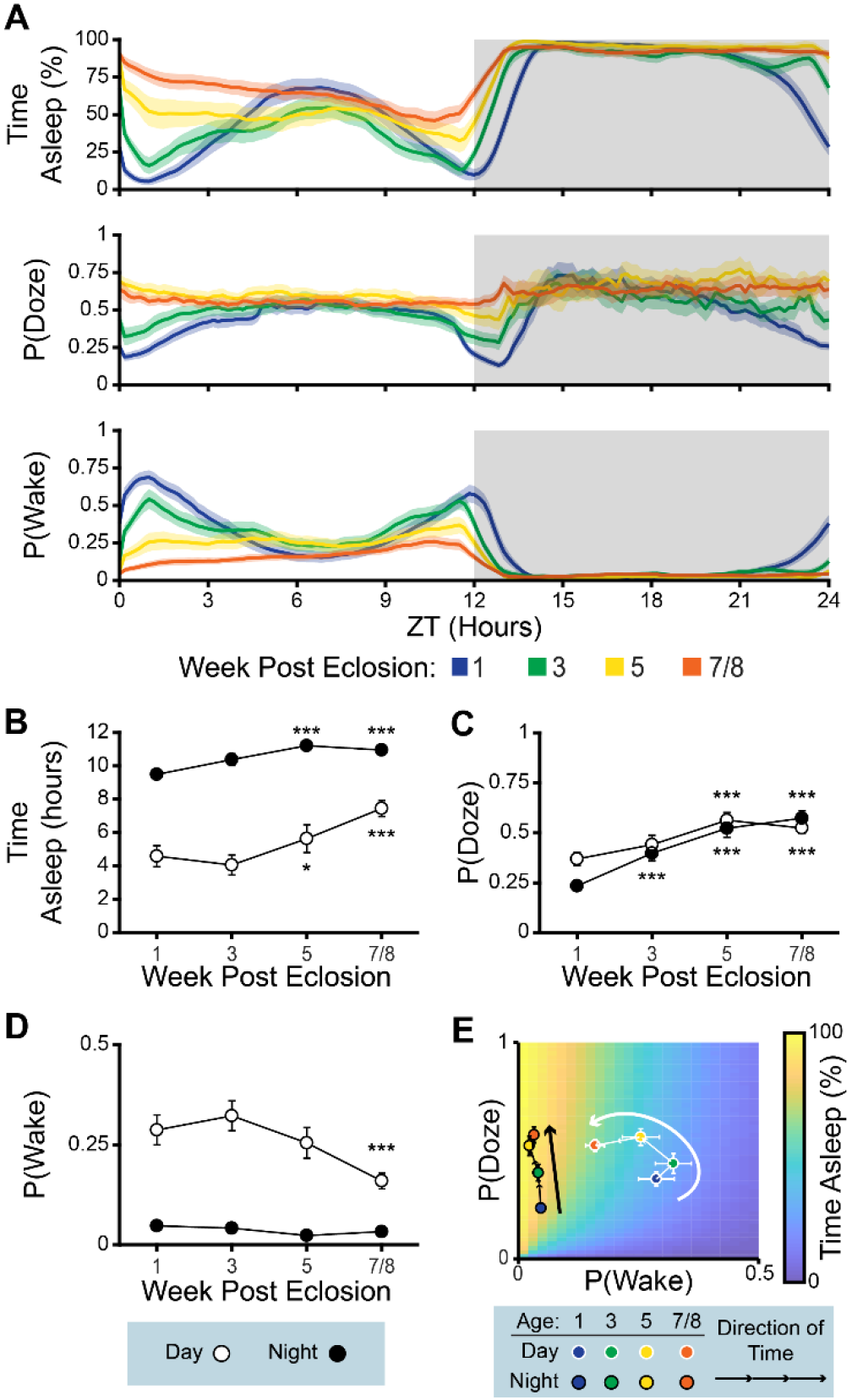
Changes in Sleep Due to Age are Due to Both Sleep Pressure and Depth. (A) Daily cycle of % time asleep, P(Doze), and P(Wake) for wildtype (CS) flies at 1, 3, 5, or 7+ weeks post-eclosion. (B-D) Comparison of time asleep (B), P(Doze) (C), and P(Wake) (D) across aging, during daytime (open circles) and nighttime (filled circles). Error bars are standard error of the mean. (E) The population means of the aging fly cohorts are overlaid on the *in silico* predicted sleep heatmap. Daytime values: white outline, Nighttime values: black outline. *n.s.* indicates no significant difference, * p < 0.05, ** p < 0.005, *** p < 0.0005.

Plotting the changes in P(Wake) and P(Doze) as a trajectory through probability space allows us to assess how aging-related changes in behavioral transition probability are related to total sleep (Fig. 5E). Nighttime behavior follows a linear path, the extent of which falls within a rather uniformly high-sleep area of the probability space, consistent with the relatively modest changes in sleep quantity at night (Fig. 5A,B). Daytime behavior follows a more complex path that moves from a low-sleep area of the space toward a high-sleep area. In this aging dataset, changes in behavior are generated by alterations in both P(Wake) and P(Doze). The effect of aging on nighttime P(Doze) is much stronger than is effect on the amount of sleep. Our method of analysis reveals an “invisible” sleep/aging interaction that would not be apparent with currently used sleep metrics. Determining the biological basis of this shift in transition probability will provide insight into the aging process.

## DISCUSSSION

Sleep is regulated by a large number of internal and external forces. Competing or interacting switches which are organized hierarchically can allow many factors to be integrated, but understanding the role of a single switch in the system can be difficult if only its effect on the amount of sleep is known. In this paper we address this fundamental problem by measuring and modeling sleep in terms of the probability of wake/sleep transitions. We define two drives, P(Wake) and P(Doze), that together can explain the amount of total sleep expressed by individual animals under a variety of conditions. Further, we demonstrate that these two drives are independently controlled and the result of different underlying biological processes.

### Probability of Wake is a Measure of Sleep Depth

The depth of sleep is clinically important and tied to its restorative power (Scammell et al. 2017). In humans, sleep depth is measured based on vital signs, EEG, and behavior, which together provide a holistic view of the process (Bonnet et al. 2007). Sleep depth in flies is more difficult to characterize. Total LFP power in flies is significantly reduced immediately after the initiation of sleep, but does not scale to indicate increased depth of sleep even as arousal threshold changes (van Alphen et al. 2013; Yap et al. 2017). Sleep depth is therefore most commonly measured in flies using arousal threshold to an applied sensory stimulus (van Alphen et al. 2013; Guo et al. 2011; Hendricks et al. 2000; Huber et al. 2004; Linford et al. 2012; Shaw et al. 2000).

Reliance on sensory arousal thresholds incurs two significant confounds: the regulation of sensory systems by time of day, and the disruptive nature of the measurement. Like other animals, the sensory systems of *Drosophila* are regulated by the circadian clock. This has been most robustly demonstrated in the response to light (Baik et al. 2018; Chen et al. 1992; Nippe et al. 2017) and to odors (Krishnan et al. 1999; Zhou et al. 2005). The major confounding feature of sensory arousal threshold, however, is that it unavoidably disrupts sleep. Rousing the flies is likely to disrupt the deeper-sleep states found in longer sleep episodes (van Alphen et al. 2013). Repeated rousing may activate homeostatic mechanisms that alter sleep depth (Hendricks et al. 2000). Finally, the arousing stimulus may act as a zeitgeber to the circadian system (Lee et al. 1996).

In light of these confounds to sensory arousal threshold, a non-invasive measure of sleep depth would be extremely helpful in understanding the function and regulation of sleep in *Drosophila*. P(Wake) is an attractive alternative measure, and several lines of evidence reinforce its validity as a measure of sleep depth. Most directly, our arousal threshold experiments demonstrate that P(Wake) correlates very significantly with ability to be aroused by either a mechanical or light stimulus (Fig. 2). We also find that the arousal-associated molecule dopamine increases P(Wake) both in chronic and acute activation experiments (Fig. 4). This is consistent with observations that the rate of spontaneous self-awakening is lower during deeper sleep stages in humans (Zepelin 1986). Considering this evidence, we propose that P(Wake) is a behavioral correlate of sleep depth in flies whose magnitude is closely aligned with dopaminergic tone.

### Probability of Doze is a Measure of Sleep Pressure

After long periods of wake, the desire to sleep also increases. This homeostasis-promoting desire to sleep is described as “sleep pressure,” or process S, in the influential Borbély two-factor model of sleep (Borbély 1982). Sleep pressure is difficult to measure directly because it is an internal state rather than an external behavior. The most common measure of sleep pressure is indirect – a change in the amount of total sleep from baseline, *i.e.* rebound sleep after deprivation (Huber et al. 2004). However, this metric is inevitably confounded by sleep depth, which increases in rebound sleep and which we show is the major driver of the amount of sleep across conditions (Fig. 1, 3, 4). Another common measure of sleep pressure is the latency to sleep after an environmental cue, such as lights turning off (Andretic and Shaw 2005; Linford et al. 2012). Sleep latency is not confounded by sleep depth, but it can only be measured relative to a pre-defined marker and only once per fly per event, limiting both its temporal resolution and statistical power.

Probability of ceasing activity, P(Doze), is conceptually similar to sleep latency. The primary difference between the two measures is that sleep latency is measured against a discrete external time marker, while P(Doze) is measured using fly-initiated activity, providing a continuous read-out of the state of the fly. Due to this similarity, we expected that P(Doze) would measure a similar biological process to sleep latency, namely sleep pressure. Empirically, we find that P(Doze) behaves as we would expect a measure of sleep pressure to behave. First, P(Doze) increases precede sleep increases and P(Doze) decreases lag behind sleep as debt is discharged (Fig. 3C, H). Second, P(Doze) is not a major determinant of total sleep in un-manipulated flies, consistent with findings showing that inhibition of the sleep homeostat has only a minimal effect on baseline sleep (Dubowy et al. 2016; Liu et al. 2016). Third, and perhaps most importantly, we show that P(Doze) has “memory trace” properties-after sleep deprivation it remains elevated until the completion of rebound sleep (Fig. 3).

While the similarities to process S are striking, the daily pattern of P(Doze) does not have the same shape as Borbély’s conceptualization (Borbély 1982). Process S is non-circadian and increases steadily during wakefulness, while P(Doze) is fairly flat with troughs during the morning and evening peaks of activity (Fig. 1) (Rieger et al. 2006). This could be due to P(Doze) not being a “pure” sleep pressure signal – it is clearly influenced by the clock and by light (Fig. 1), tying P(Doze) to the animals’ internal and external states. This is consistent with the idea that the homeostat has a relatively minor role in the specification of sleep drive in unperturbed animals, and is only engaged in the case of deviation from the norm. Notably, in our sleep deprivation experiments the trajectory of P(Doze) becomes much more similar to S as sleep debt builds up (Fig. 3C and H). Alternatively, it may be that the two-process model simply does not reflect regulation of sleep pressure in *Drosophila*. Recent evidence suggests that sleep pressure in flies is primarily regulated by the clock rather than by sleep debt (Geissmann et al. 2019). The lack of agreement between model and empirical results suggests the need for a revised and more broadly-based model of sleep drives.

Counter-intuitively for a measure of sleep pressure, P(Doze) is correlated with increased sleep fragmentation (Fig. 5). P(Doze) is the probability of initiating sleep episodes, and since the median sleep episode is shorter than the mean episode in the typical distribution of episode durations (Diamond 2016; Ueno et al. 2012a), it follows that increasing P(Doze) will usually result in shorter mean sleep episode duration. The important implication of this finding is that sleep fragmentation is not, on its own, evidence of a wake-promoting process. Our results indicate that fragmented sleep can be the result of decreased sleep depth (*i.e.* high P(Wake)), increased sleep pressure (*i.e.* high P(Doze)), or a combination of the two. Biologically this may make sense. Allowing fragmentation of sleep to be generated by a process untethered to arousal threshold or dopaminergic tone gives the brain more degrees of freedom in determining sleep pattern.

### Independence and Integration of P(Wake) and P(Doze)

Our analytical method defines two parameters that can assess the core drives that determine the sleep state. Importantly, these parameters are independent - we show that P(Doze) and P(Wake) are only moderately correlated with one another. This dissociability is quite clear in the response of these parameters to different perturbations. Increases in dopaminergic tone have selective effects on P(Wake), while dietary sucrose, which affects sleep structure, acts via changes in P(Doze).

In contrast, other perturbations of the system (*e.g.* aging, the *insomniac* mutation, and mechanical sleep deprivation) alter both P(Doze) and P(Wake). During sleep deprivation P(Doze) measures sleep need, but decreased P(Wake) is the primary driver of the amount of rebound sleep (Fig. 3), indicating that the two processes can also interact. These interactions may give additional insight into the privileged roles of some neural circuits in increasing total sleep after sleep deprivation (Seidner et al. 2015) and in the into the ability of some forms of sleep deprivation to bypass engaging the homeostat (Beckwith et al. 2017).

### Conclusions

Classifying behavior into discrete states to determine transition probabilities and pathways is an approach that has been applied to many behaviors, including swimming (Willows and Hoyle 1969), grooming (Berridge et al. 1987), feeding (Martaresche et al. 2000), and navigation (Hernandez-Nunez et al. 2015; Kato et al. 2015). In this report, we treat sleep and wake as binary behavioral states and measure the probability that animals transition between them. As our understanding of sleep grows, it has become clear that two individuals displaying the same amount of sleep may have arrived at that amount on very different paths. The ability to characterize a sleep set point in terms of the animal’s underlying arousal state and sleep drive will help tease apart the complexities of the circuits that specify sleep.

## METHODS

### Animals

Flies were raised on cornmeal-dextrose-yeast food in a 25°C incubator. The incubator maintained a 12 hour:12 hour light:dark cycle. Flies were collected 0-2 days post-eclosion and housed in mixed-sex, low-density vials until the beginning of the experiment (typically 12 male and 12 female flies). Female flies used in behavioral experiments were co-housed with males and were therefore presumably mated. For experiments in which the age of the flies is not specified, flies were 5 – 10 days old at the beginning of data collection. Flies used in the aging experiment were housed in large fly culture bottles with cornmeal-molasses food in order to facilitate a large population surviving to the 7/8 week time point. Transgenic flies were as follows: *per*^*01*^(CS) – gift of Michael Rosbash, *w*^*1118*^ and *w*^*1118*^; *dat*^*fmn*^ – gift of Amita Sehgal, PBac{WH}*inc*^*f00285*^, *w*^*1118*^ (*inc*^2^; BDSC # 18307), *TH-Gal4* (BDSC # 8848), *UAS-dTrpA* (Hamada et al. 2008).

### Sleep Data Collection

For DAM system experiments, flies were briefly anesthetized with CO_2_, loaded into glass tubes containing 5% agarose/2% sucrose food (unless otherwise specified), and tubes were mounted in *Drosophila* Activity Monitor (DAM) boards (Trikinetics, Waltham, MA). Experiments were carried out at 25°C with a 12 hour:12 hour light:dark cycle, unless otherwise specified. Flies were given a day to acclimate to the DAM system before data collection began. Sleep deprivation was performed by securing the DAM systems to a motorized vortexer and shaking the flies with an inter-pulse period of 10 seconds and a pulse duration of 2 seconds. Unshaken control flies were housed in the same incubator, but not attached to the vortexer.

For Flybox experiments (Guo et al. 2016), flies were briefly anesthetized in a 4°C cold room and loaded individually into a 96-well plate, each well containing 300 µL of 5% agarose/2% sucrose food. Plates were securely fitted into Flyboxes. Images were acquired at 0.1 Hz, and flies were tracked in real time using custom Matlab code. Experiments were carried out with a 12 hour:12 hour light:dark cycle and flies were given a day to acclimate to the Flybox before data collection began.

### Arousal Threshold Measurements

Light-driven arousal threshold in the DAM system was measured in a dimly lit incubator (40 lux daytime) by delivering 3 second pulses of red light (approximately 250 lux) at the times specified. Mechanical arousal threshold in the Flybox was measured by applying a single tap to the side of the 96-well plate using a solenoid (Guo et al. 2016) at the times specified. Stimulated arousal rate was calculated as the fraction of flies that were previously asleep that became active during the minute following the stimulus. Unstimulated arousal rate was calculated by performing the same calculation at a timepoint 6 minutes before the delivery of the stimulus. The Fisher’s Exact test was used to determine if the rate of awakenings following a stimulus was significantly different from the spontaneous rate of awakenings. % Response to tapping is the difference between the stimulated and unstimulated arousal rate, *i.e.* the excess awakenings beyond the spontaneous rate.

### Generation of In Silico Data

Simulation data was generated by custom programs written in Matlab (MathWorks, Natick, MA). The simulation accepted as input the number of simulated flies to generate, the duration (in time steps) of each simulation, and the values of P(Wake) and P(Doze) for the *in silico* flies. The simulation was initialized in a random state with equal probability of wake and sleep at *time* = 1. At each subsequent time step *i*, the simulator: 1) Generated a random number between 0 – 1, 2) Checked activity at *time*(*i* – 1), 3) Compared the random number with the behavioral transition probability. If the fly was active at *time*(*i* – 1), it was compared with P(Doze). If the fly was inactive, it is compared with P(Wake). 4) If the random number was less than or equal to the appropriate transition probability, the fly switched states. 5) The simulation proceeded to the next time step. The first 100 time steps were discarded to minimize the contribution of the initialization state to the simulation output. Activity was coded as 1, inactivity was coded as 0.

### Sleep Data Analysis

DAM system, Flybox, and *in silico* data were analyzed with a common set of modular programs written in Matlab. Raw beam crossing data from the DAM was trimmed to the appropriate experimental time period and binned to 1 minute intervals using DAMFileScan (TriKinetics). Flybox tracking data was converted to distance traveled per frame, thresholded at 10 pixels (approximately 1mm), and binned to 1 minute intervals. The experimental design and fly genotype metadata were transcribed into a text file. The Matlab program imported the movement data and metadata. Flies with 13 or more hours of continuous inactivity at the end of an experiment were excluded as potentially dead. Continuous periods of inactivity lasting 5 minutes or longer were classified as sleep (Shaw et al. 2000).

Probability of wake, or P(Wake), was calculated as follows: 1) Count every bin of inactivity except the last one. This is the denominator, the total number of state transitions from inactivity. 2) For every bin of inactivity, count the number of times the fly is active in the subsequent bin. This is the numerator, the total number of state transitions from inactive to active. 3) Divide the total state transitions from inactive to active by the total state transitions from inactive. If there were no bins where the fly does not move (*i.e.*, denominator is zero), the transition probability is undefined. Probability of Doze, or P(Doze), was calculated identically, but with activity and inactivity reversed.

Statistical analysis was performed using the Matlab statistics toolbox, data was tested for main effects using ANOVA, and post-hoc t-tests were corrected using Tukey’s Honestly Significant Difference Procedure (hsd). Plots of sleep measures across time of day were calculated using a moving average with width of 30-90 minutes at 10 minute intervals.

## ACKNOWLEDGEMENTS

We would like to thank Dr. Jané Kondev and the students in the Brandeis Quantitative Biology Research Community (QBReC) who assisted in collecting preliminary data, and Dr. Michael Rosbash and the Brandeis Science Communication Lab for helpful comments on this manuscript.

## SUPPLEMENTAL STATISTICAL TABLES

**Figure S1.**
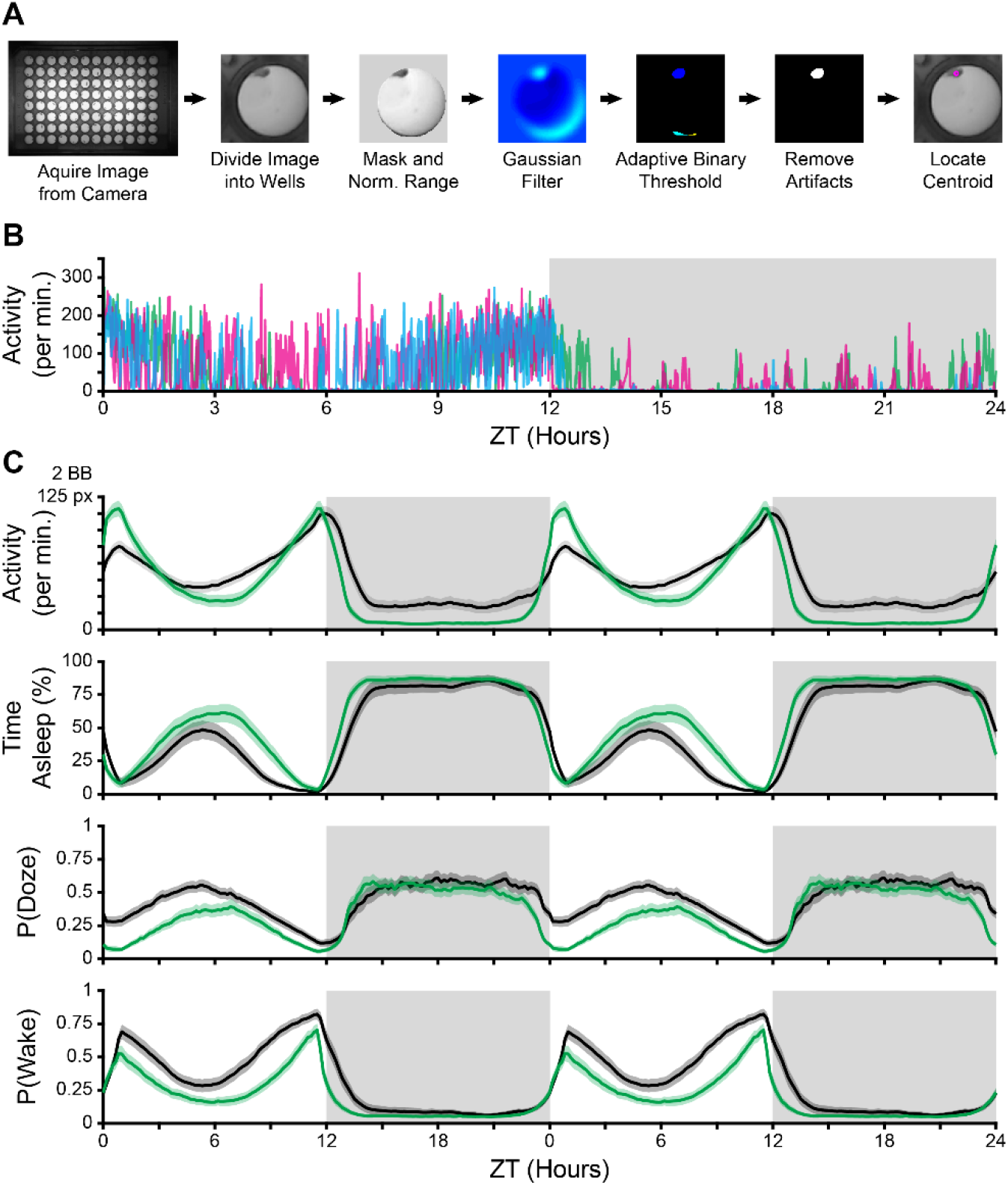
Video Tracking of Activity and Sleep in the FlyBox. (A) Analysis pipeline for locating the coordinates of each fly in each frame. (B) Activity, measured in pixels traveled per minute, plotted for 3 individual flies (cyan, green &magenta). (C) Activity, time asleep, P(Doze), and P(Wake) are plotted for CS flies in the DAM system (Black lines, reproduced from Figure 1), and in the FlyBox (Green lines, *n* = 55). Curves are the double-plotted average of 3 days of activity. BB: beam breaks, px: pixels. Gray area indicates lights-off period.

**Figure 1:**
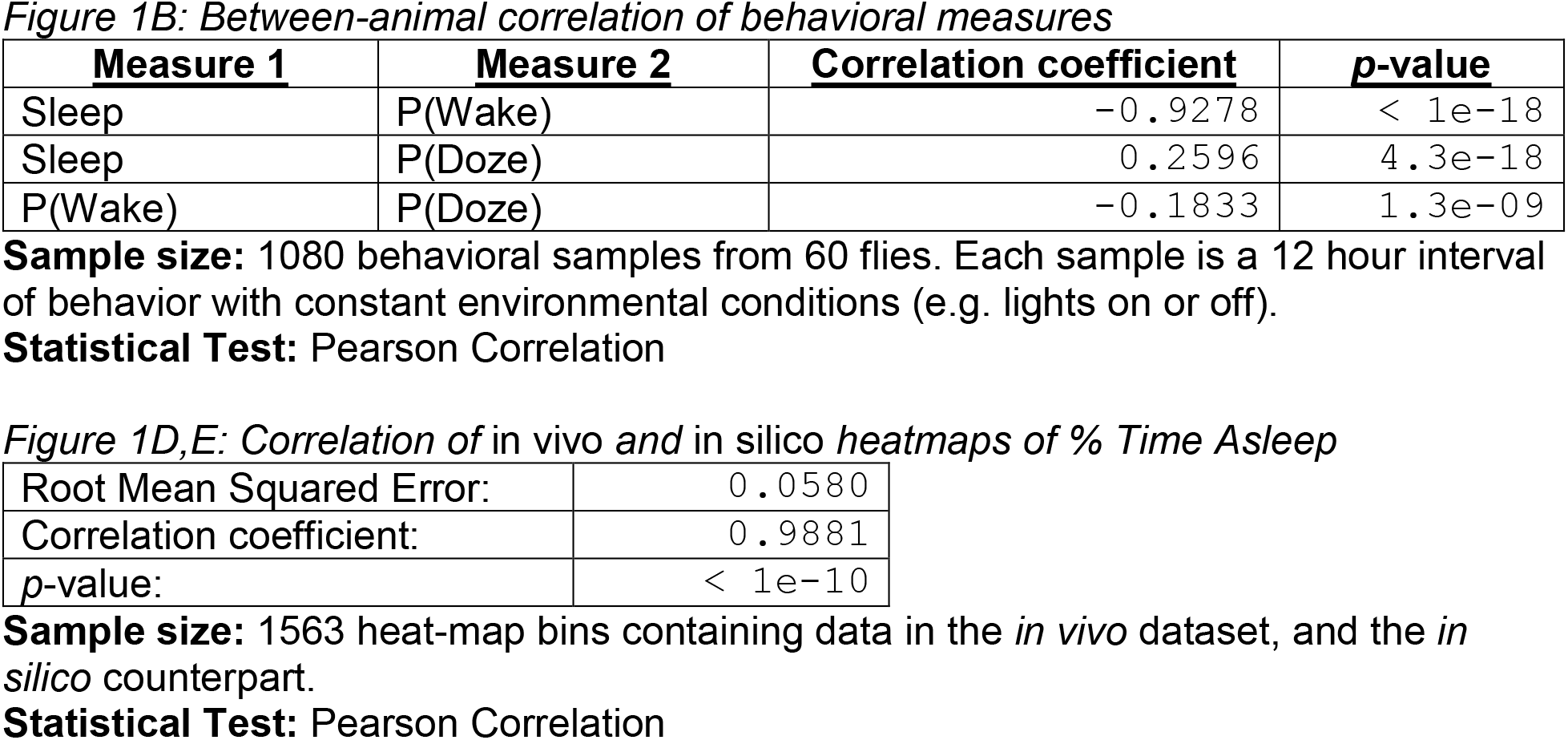

**Figure 2:**
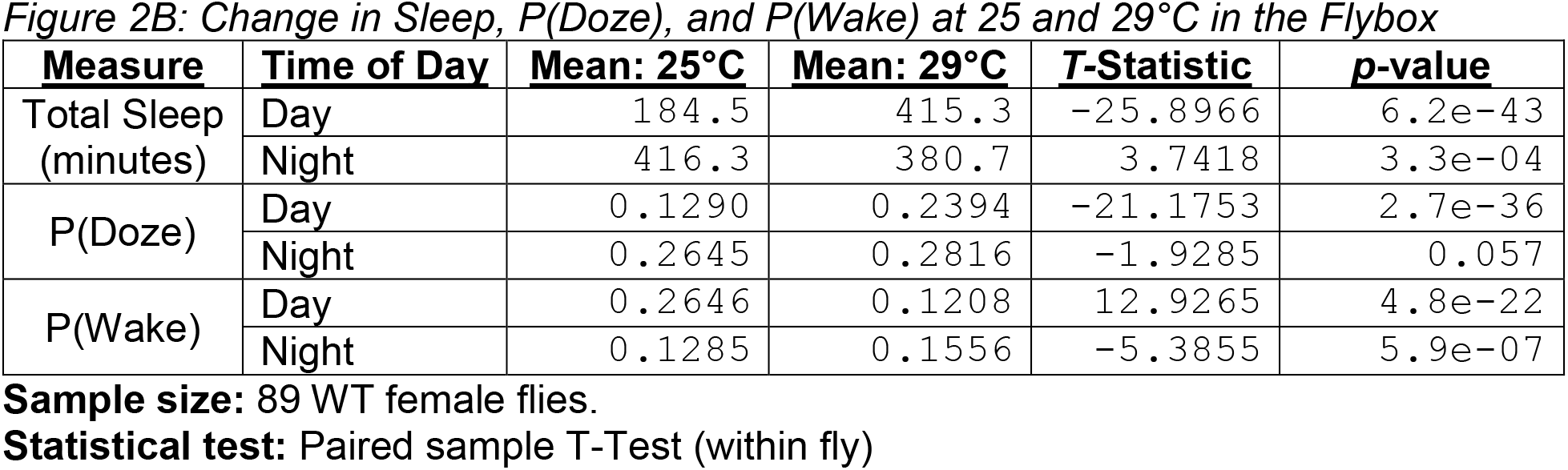
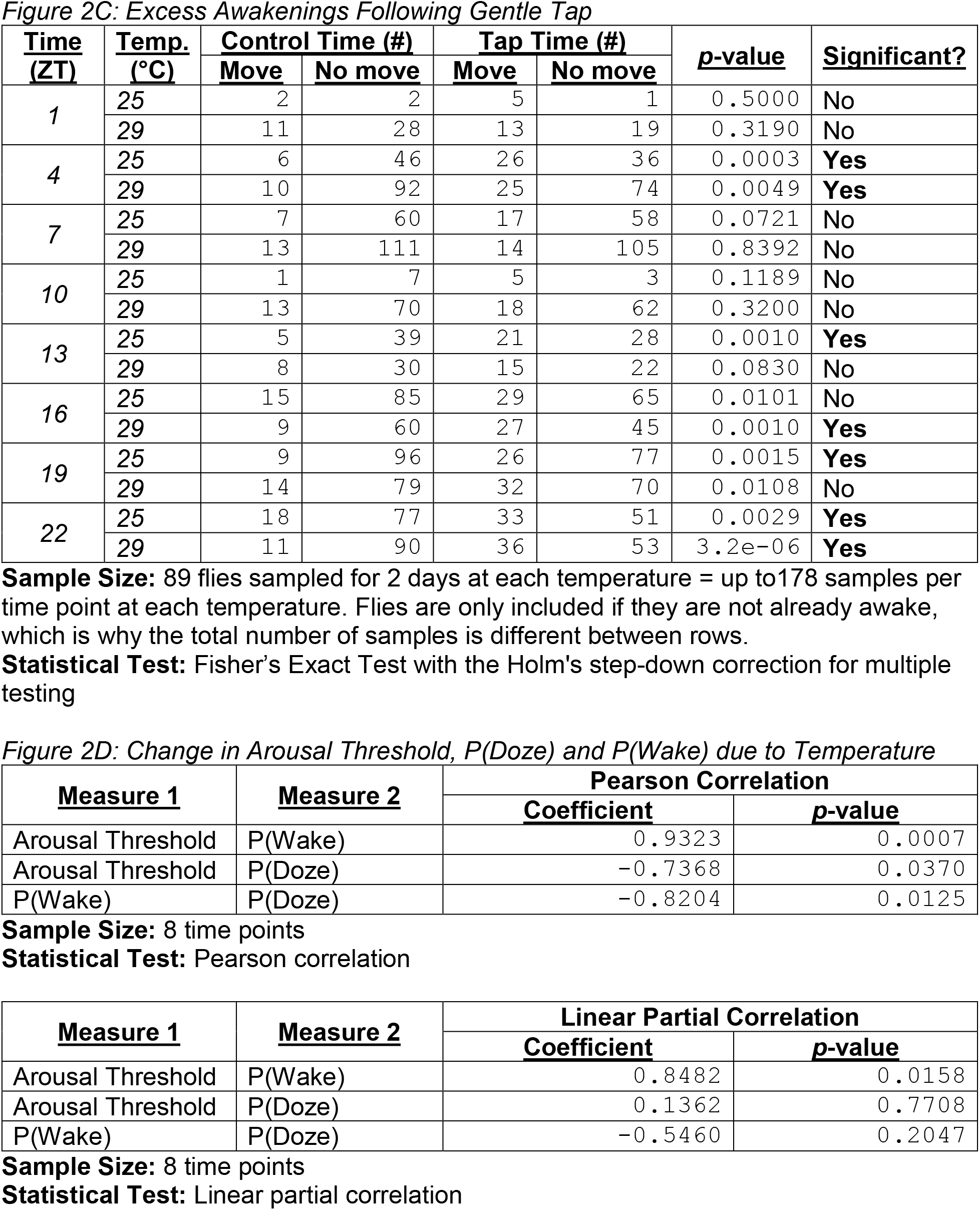
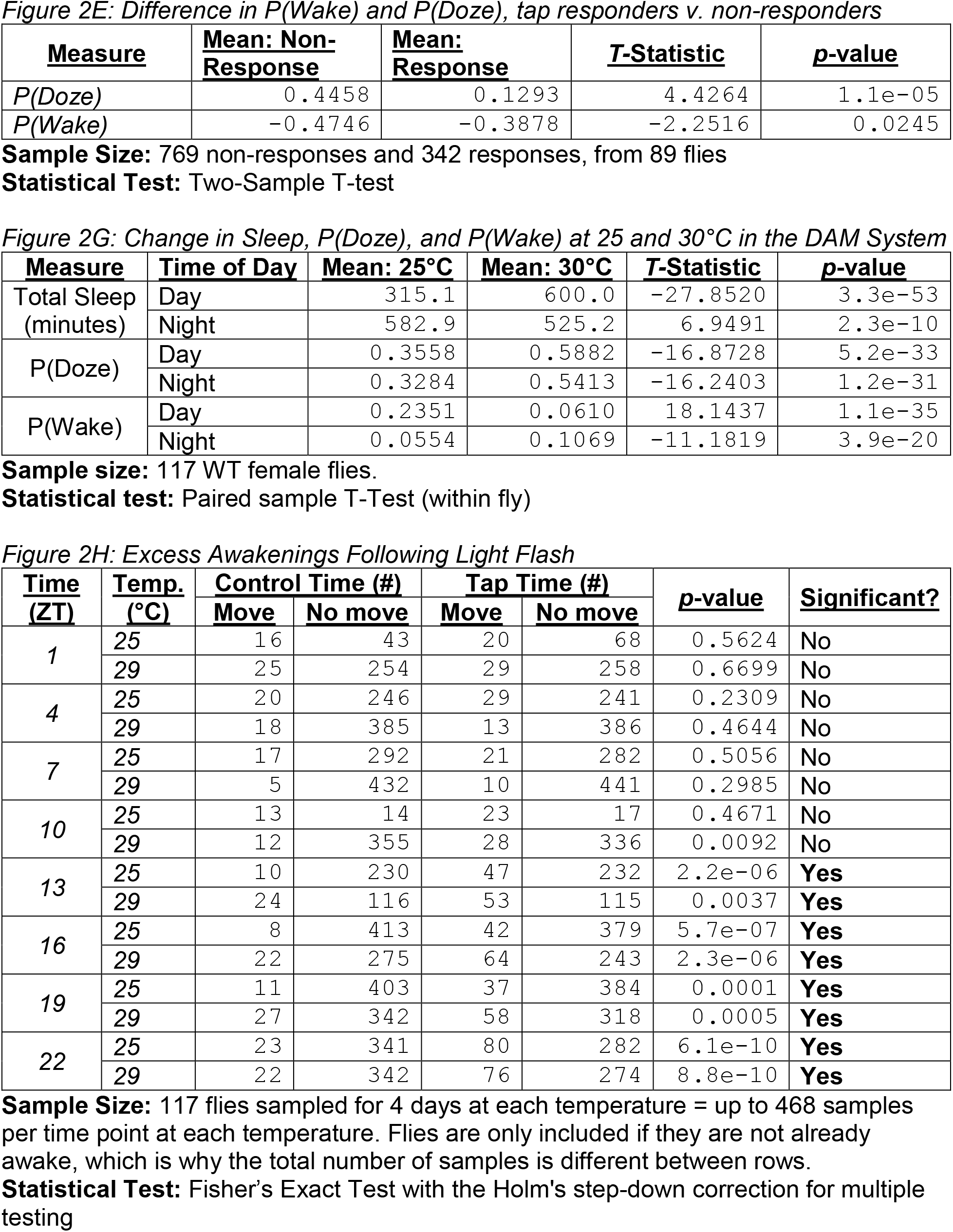
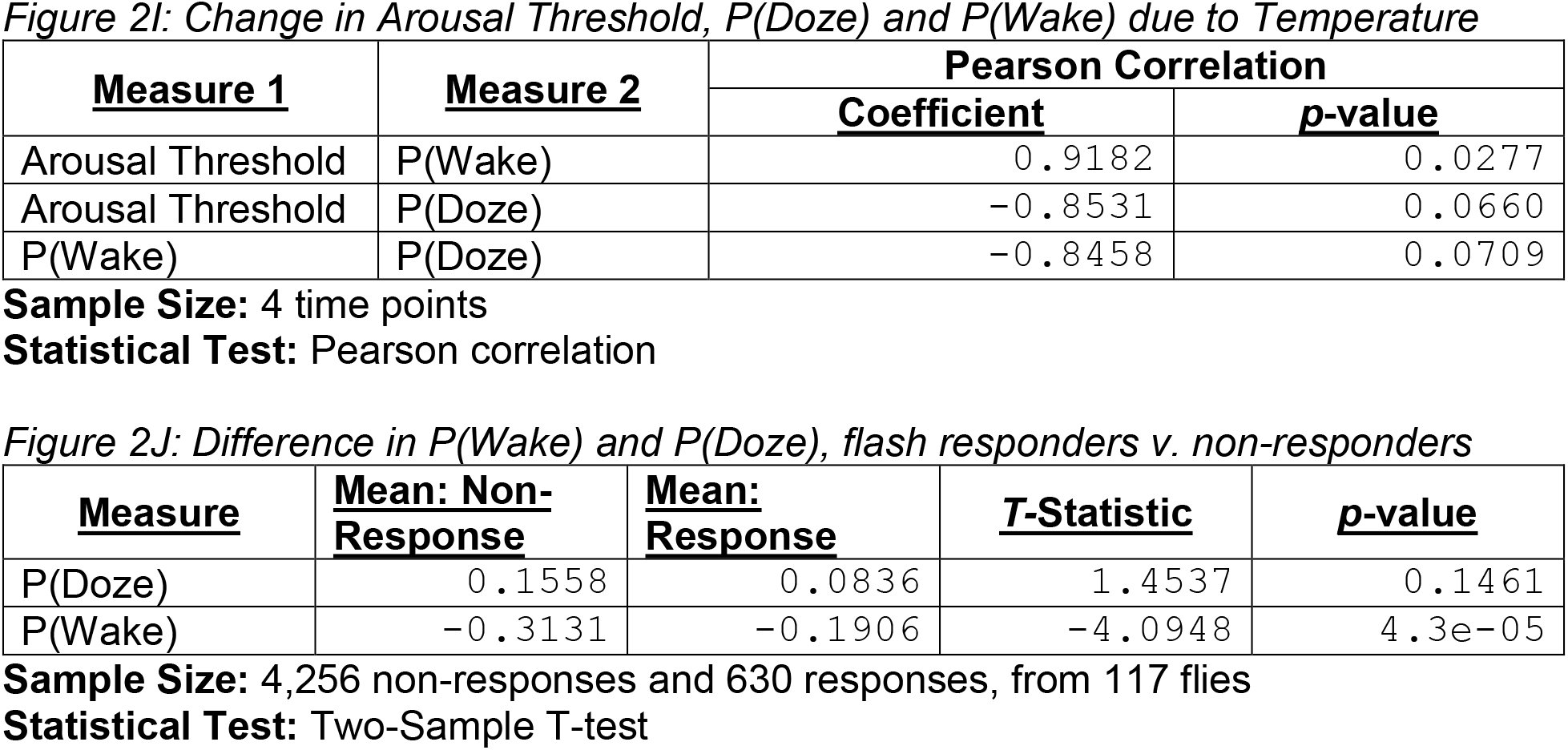

**Figure 3:**
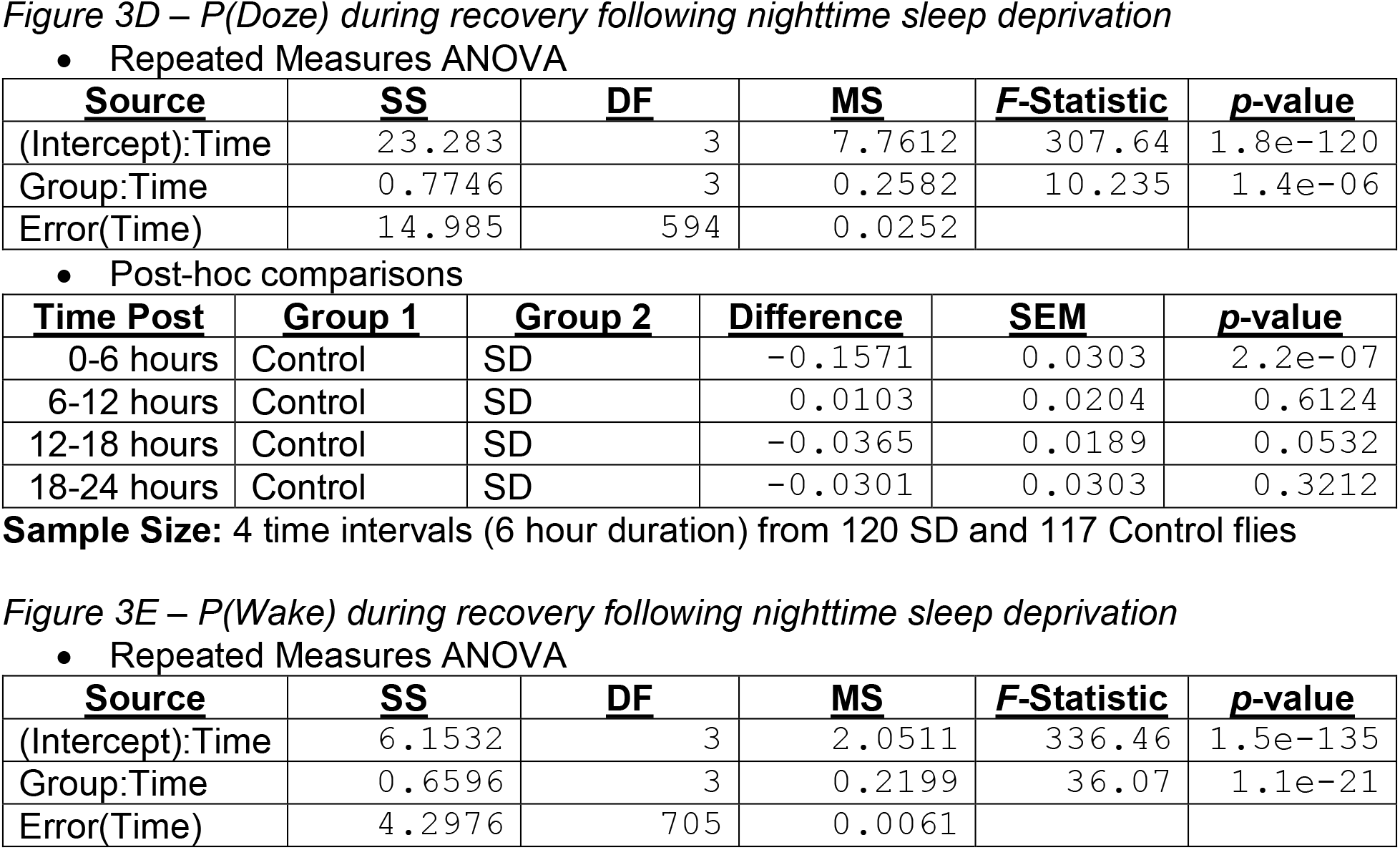
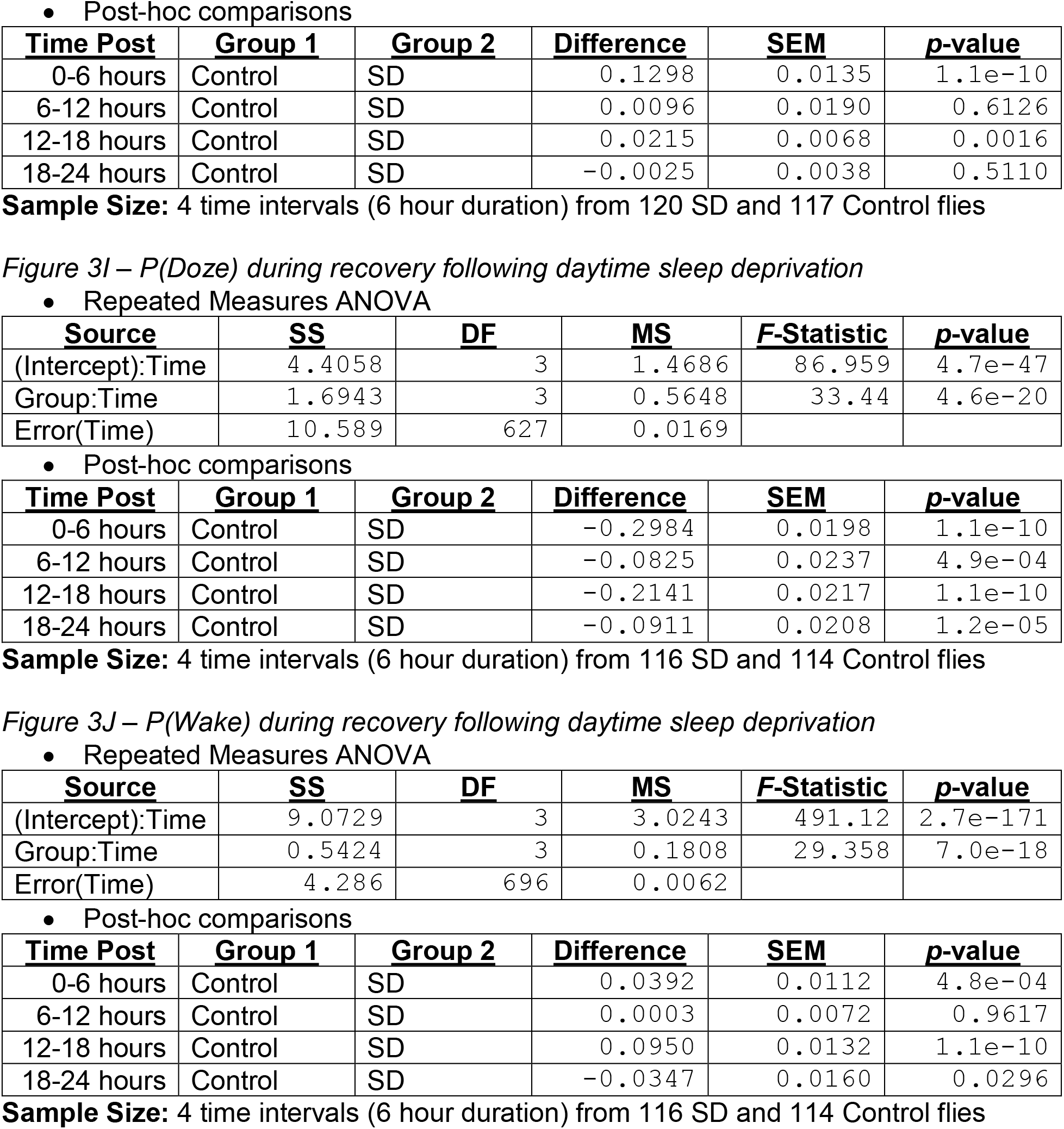

**Figure 4:**
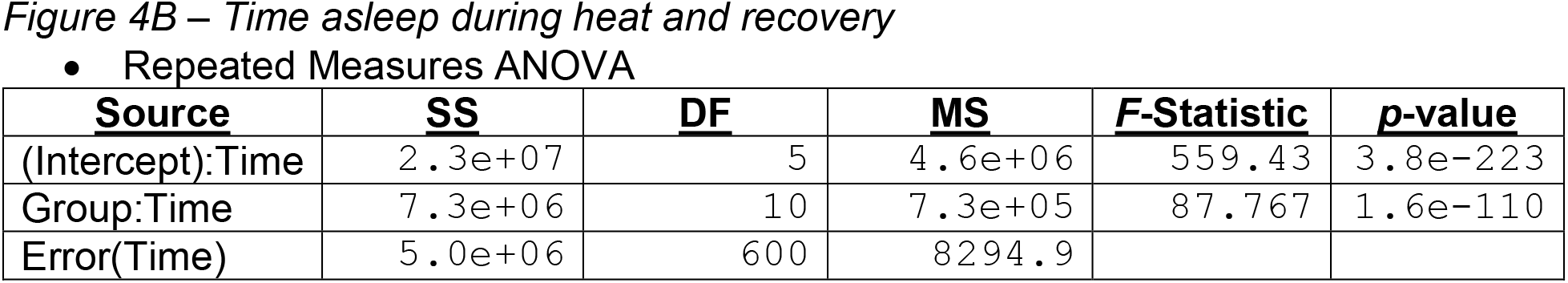
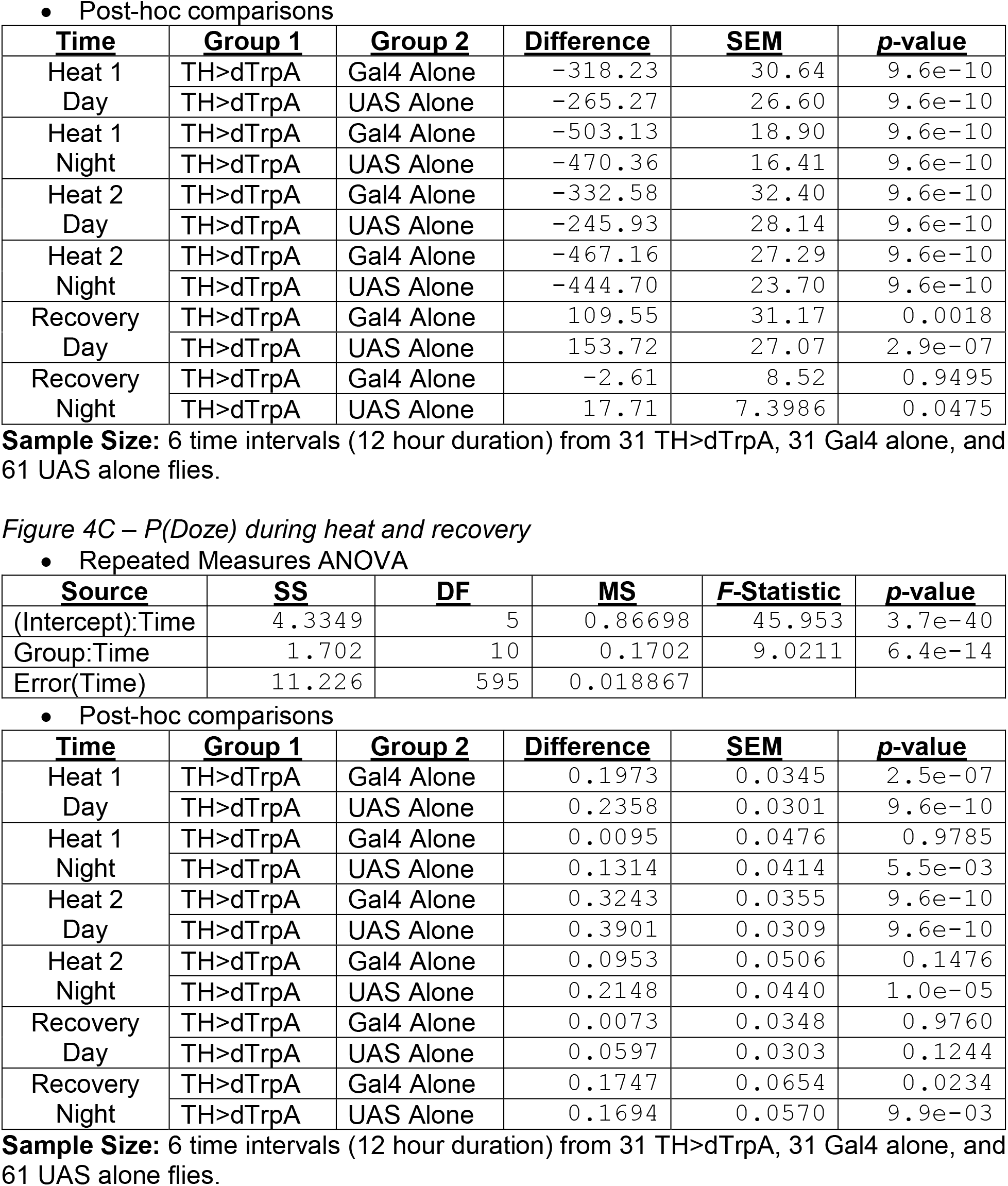
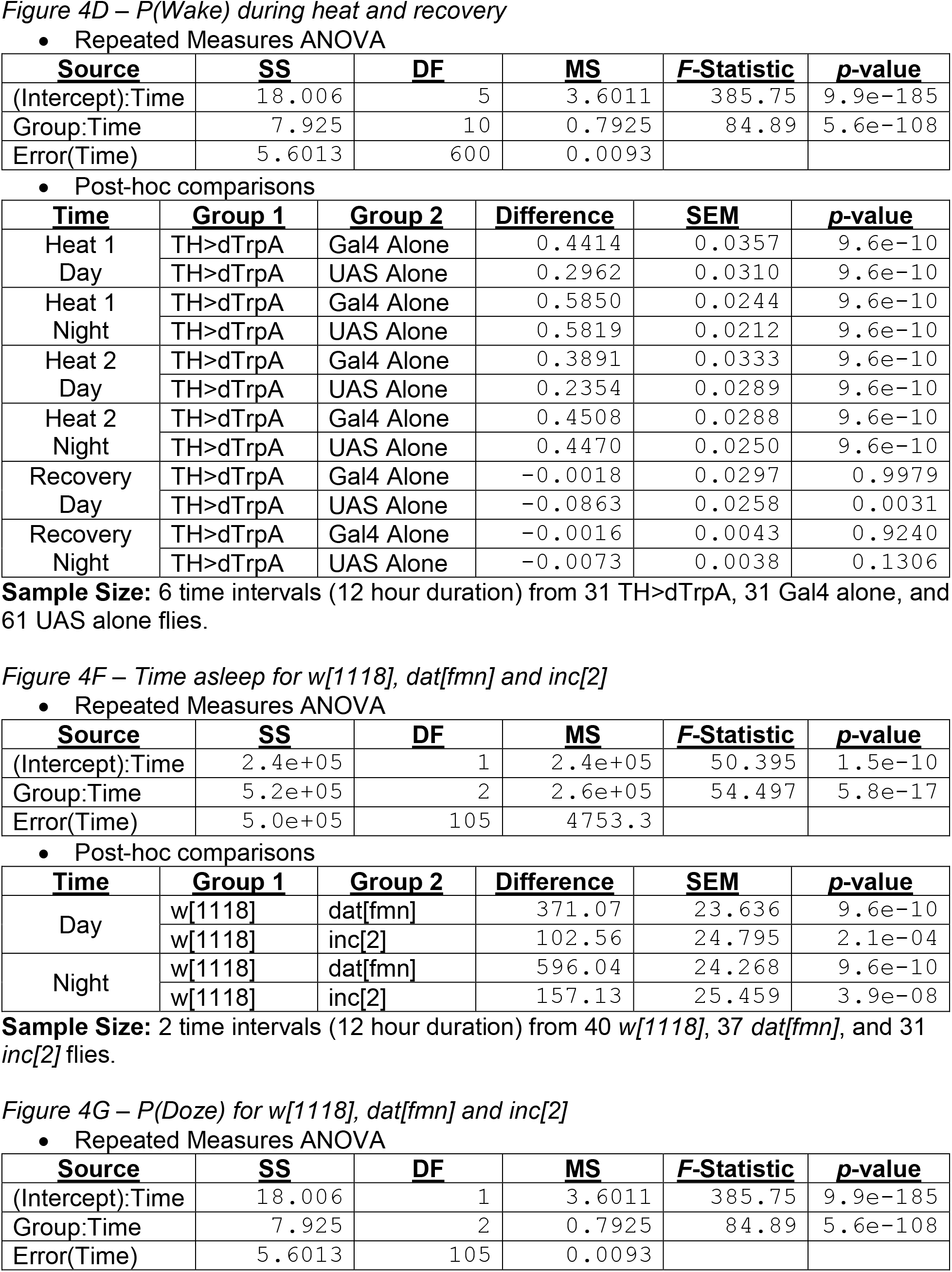
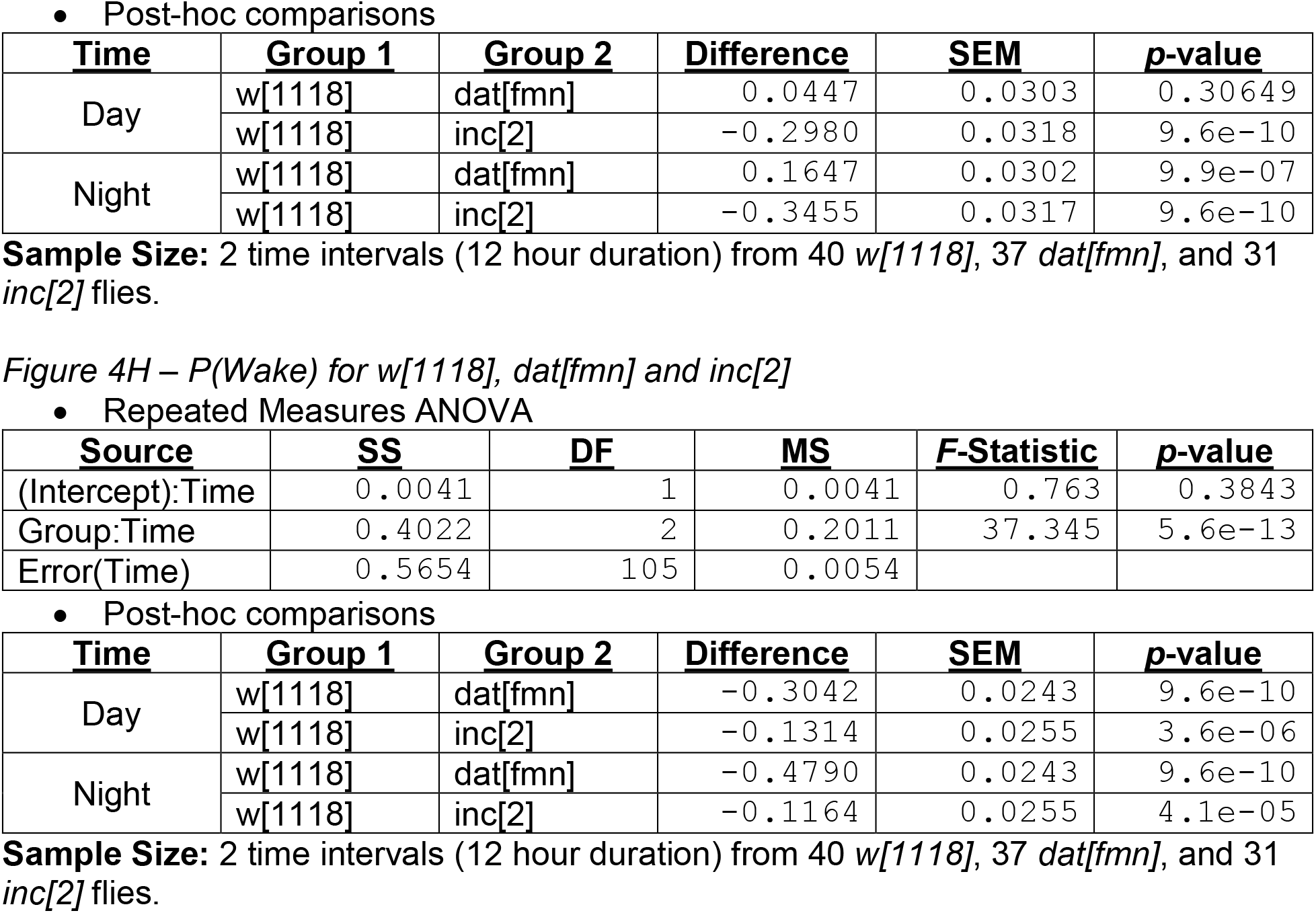

**Figure 5:**
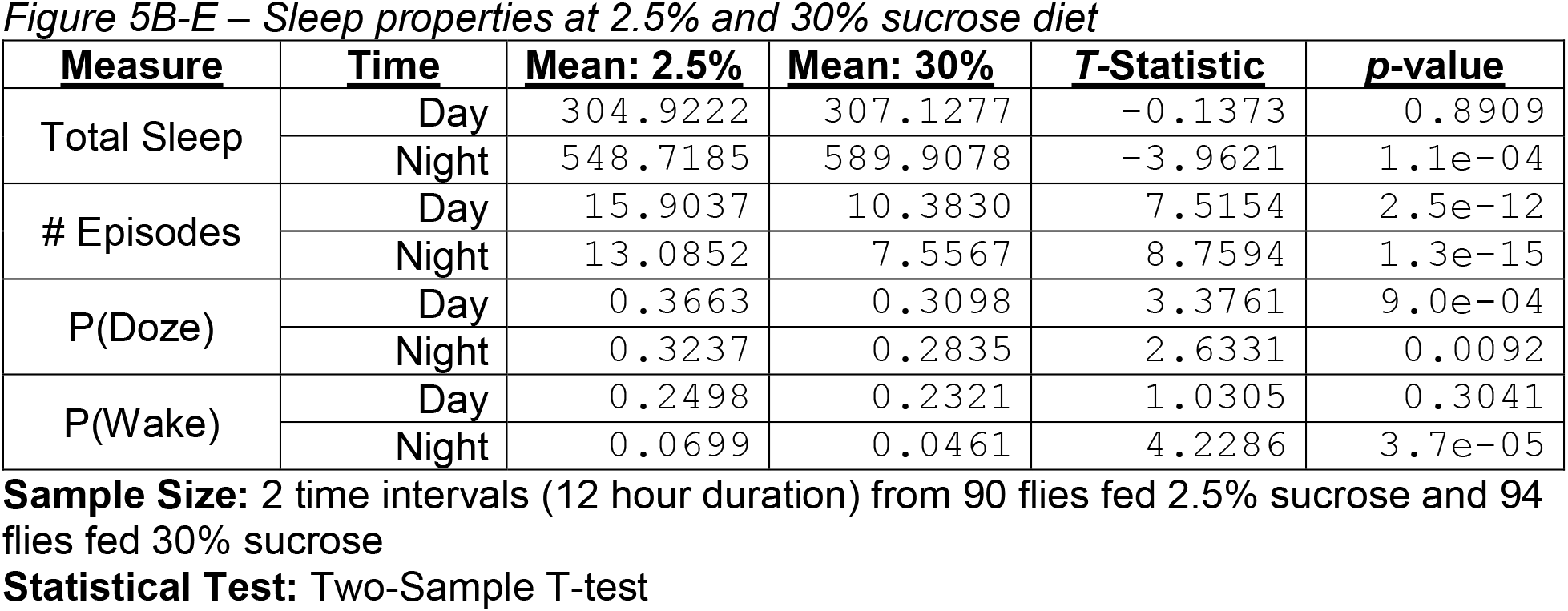

**Figure 6:**
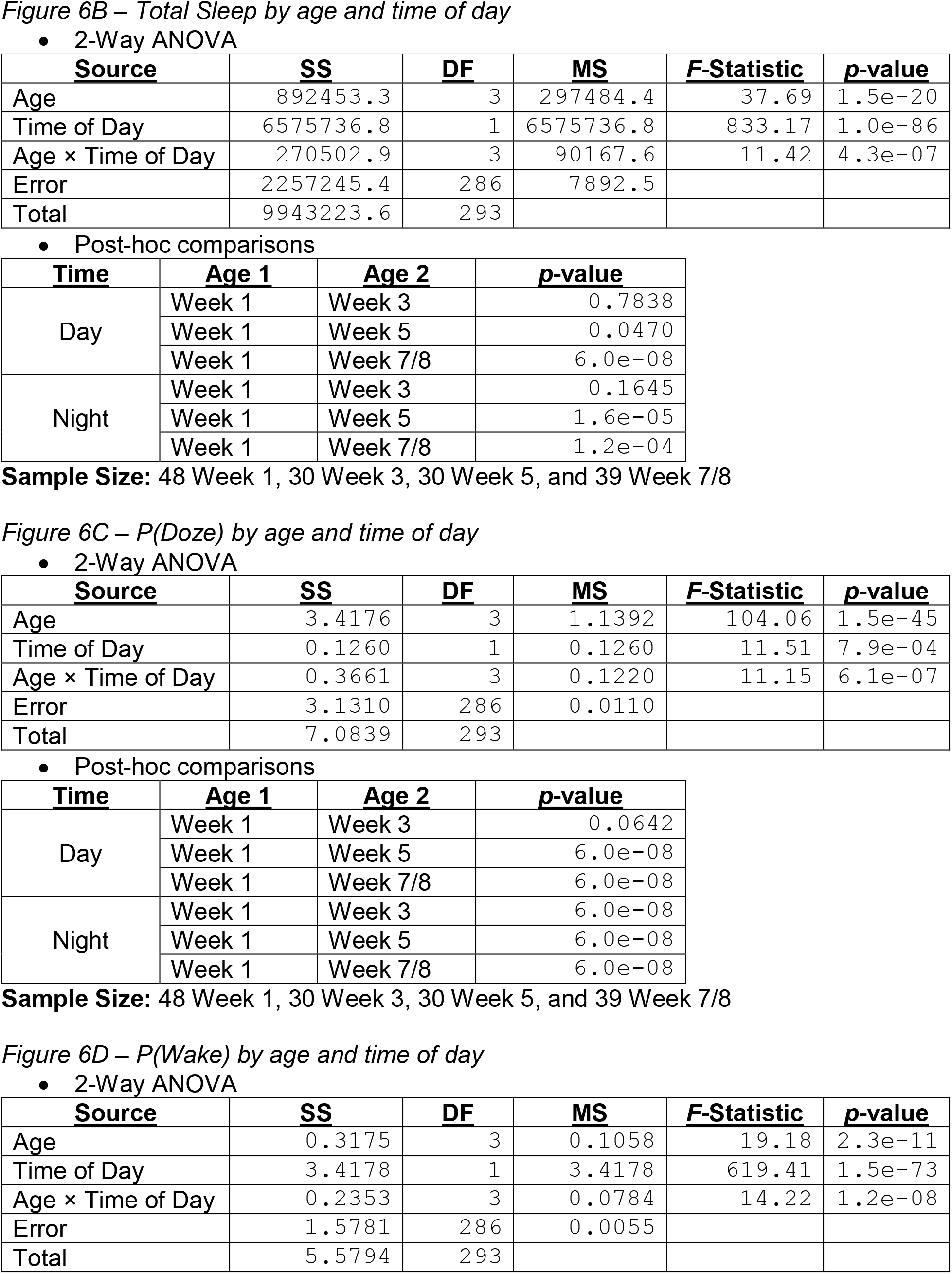
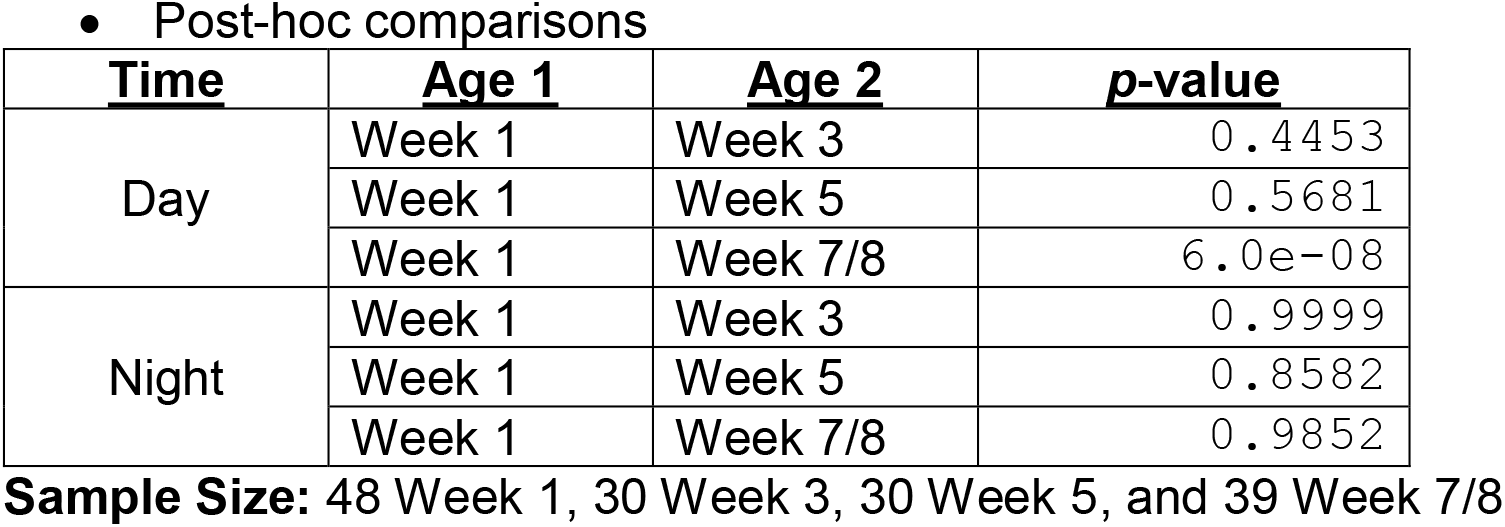

